# Simulating protein evolution with programmable fitness objectives derived from modern protein models

**DOI:** 10.64898/2026.10.26.761958

**Authors:** Kaiyu Qiu, Andrei N. Lupas, Stanislaw Dunin-Horkawicz

## Abstract

Simulation of protein sequence evolution provides a controlled way to test whether specified evolutionary mechanisms can generate the patterns observed in natural proteins. However, classical simulation largely relies on fixed substitution rules, whereas existing fitness-based simulations often define fitness using a single proxy and thus do not sufficiently capture the complex nature of constraints imposed by selection. Here, we simulate protein evolution through amino acid substitutions under programmable fitness objectives derived from individual or combined protein models, including protein language models, inverse folding models, physics-based energy functions, and alignment-based statistical models. Evaluation of simulations revealed that different fitness proxies impose distinct trade-offs between evolutionary realism and molecular plausibility, motivating the use of combined objectives to redistribute selective pressure across complementary constraints. We then show that programmable fitness objectives can encode mechanism-specific biological contexts. Incorporating oligomeric structural context recovered interface-specific evolutionary constraints in homodimeric proteins, while multi-conformation objectives better reproduced natural sequence variation in metamorphic proteins and provided interpretable estimates of how alternative native folds contribute to evolutionary constraint. Finally, we explore a differentiable extension of this framework, in which structural objectives are backpropagated through AlphaFold2 to guide sequence evolution. Under this formulation, evolved sequences showed favorable shifts in several molecular properties relative to those from discrete, stepwise mutation-selection simulations while retaining key evolutionary signatures.

## Introduction

Sequence evolution simulation has long served as a bridge between mechanistic models and observed biological diversity. By generating artificial evolutionary trajectories under specified assumptions, simulation provides a controlled way to ask whether an assumed evolutionary mechanism is sufficient to produce patterns observed in natural sequences. This makes simulation complementary to phylogenetic reconstruction: whereas phylogenetic reconstruction infers evolutionary history from extant sequences, simulation starts from explicit evolutionary rules and examines their consequences. As a result, sequence simulation has been widely used for benchmarking phylogenetic reconstruction (1–3), hypothesis testing (4–6), parameter estimation (7), generation of realistic synthetic sequences for training machine learning (8–11), and even forecasting evolutionary dynamics (12). Protein evolution is strongly constrained by the need to maintain structure and function (13), making simulation a natural way to explore how selective constraints shape movement through protein sequence space.

Sequence simulation tools account for a wide range of evolutionary events, including substitutions, insertions, deletions, and higher-level processes such as motif rearrangement or domain shuffling (14–16). Among these, substitution is the most fundamental and is therefore considered by virtually all simulation tools. The standard approaches model substitutions as continuous-time Markov processes, applied either to forward trajectories or along a given phylogeny. Empirical substitution models such as JTT (17), WAG (18), and LG (19) define 20×20 amino acid exchangeability matrices, and simulation tools like INDELible (20), AliSim (21), and Pyvolve (22) rely on these models to introduce mutations.

Substitution-focused simulation tools typically rely on simplifying assumptions that limit their ability to represent realistic constraints. For example, the common assumption of site independence neglects epistatic interactions that are critical for protein structure and function (15). More generally, these models lack an explicit consideration of fitness and sample mutations without explicitly evaluating their functional consequences (23). As a result, key features of natural evolution are therefore approximated indirectly. For example, the evolutionary rate heterogeneity across protein sites is often modeled using Gamma-distributed rates or fixed fractions of invariable sites. More flexibly, covarion models allow site-specific rates to change across lineages, as implemented for example in MrBayes (24, 25), while other methods infer site-specific rates from multiple sequence alignments (MSAs) and incorporate them as priors (15, 21). These approaches, however, lack an explicit molecular explanation for site-specific rate variation (26) and thus may produce trajectories that deviate from natural evolution in their statistical and biophysical properties, undermining their utility in downstream tasks that require realistic variation. This limitation has been underscored by recent work showing that machine learning models can easily distinguish natural MSAs from simulated ones (26).

In contrast to sequence simulation methods that do not consider fitness effects, natural evolution is shaped by selection acting on the fitness effects of mutations, with beneficial variants generally favored and deleterious variants selected against. Accurately modeling this process requires evaluation of mutation effects, which has progressed from early methods based on sequence conservation (e.g., PolyPhen (27)) or thermodynamic stability (e.g., Rosetta (28)) to learning-based frameworks such as alignment-dependent statistical models (e.g., direct coupling analysis, DCA (29, 30)) and deep protein models enabling zero-shot prediction (e.g., protein language models, pLMs (31)). These developments enable increasingly informative proxies for protein fitness and provide an opportunity to simulate protein evolution under explicit fitness constraints, thereby bringing sequence simulation closer to the mutation– selection process that shapes natural proteins.

The application of such proxies in fitness-aware simulations has been explored for over two decades. For example, Parisi et al. modeled protein structures using contact matrices and estimated mutant energies via empirical potentials, revealing fold-dependent substitution patterns (32). Arenas et al. incorporated misfolding effects to better capture stability constraints using a contact-based free energy function (23), while more recent work has leveraged all-atom energy functions from Rosetta to simulate evolution (33). Another route to fitness-constrained simulation comes from statistical models inferred from natural alignments. In particular, DCA-based models define family-specific fitness landscapes that can drive iterative mutation–selection procedures in both discrete and continuous evolutionary frameworks (34–38). These approaches have been applied to study natural or experimentally driven evolutionary dynamics, including the emergence of epistatic signals, the reproduction of natural phylogenetic patterns, and the modeling of directed evolution experiments (23, 36, 38).

Despite these advances, the fitness function in most existing methods is defined by a single objective such as a physical energy or another learned proxy, and the resulting trajectories are evaluated for their resemblance to natural patterns. This strategy has been powerful, but it has left three aspects less explored: (i) how recent protein models perform as fitness proxies for sequence simulation, (ii) whether multiple fitness objectives can be flexibly combined to capture complementary constraints, and (iii) whether fitness objectives can be programmed to represent specific biological mechanisms beyond those imposed by monomeric globular structures (e.g. oligomeric assemblies or metamorphic proteins adopting multiple native conformations). A recent approach that bypasses the need for an explicit fitness proxy is exemplified by the PEINT (39) and DPLM-Evo (40) frameworks, which learn evolutionary sequence transformations directly from data. While capable of generating realistic evolutionary trajectories, such models rely on extensive training data and lack an explicit, interpretable fitness objective that can be directly modified to represent a hypothesized constraint. For example, one may want to ask whether oligomeric interfaces impose detectable site-specific constraints, whether multiple native conformations jointly shape the evolution of metamorphic proteins, or how sequence-level naturalness and structure-level compatibility should be balanced within the same evolutionary process. These diverse scenarios can be addressed in simulation using fitness objectives that can be flexibly constructed, combined, and interpreted, allowing protein sequence simulation to test explicit hypotheses about the molecular constraints that shape protein evolution.

Here, we simulate protein evolution through amino acid substitutions under programmable fitness objectives derived from protein language models, inverse folding models, physics-based energy functions, and/or alignment-based statistical models. We first establish an evaluation framework for comparing these objectives by examining whether simulated trajectories recover multiple signatures of natural evolution, including site-specific amino acid distributions, evolutionary rate heterogeneity, substitution preferences, and residue–residue co-variation. We then show that different objectives impose distinct trade-offs between evolutionary and molecular realism, motivating composite objectives that redistribute selective pressure across complementary constraints. Moreover, we demonstrate that programmable fitness objectives can encode specific biological mechanisms. Incorporating oligomeric structural context recovers interface-specific evolutionary constraints, while multi-conformation objectives better reproduce natural sequence variation in metamorphic proteins and provide interpretable estimates of how alternative native folds contribute to evolutionary constraint. Finally, we explore how this principle can be extended to differentiable structural constraints through AlphaFold backpropagation, showing that stronger structure-preserving objectives can improve the molecular properties of evolved sequences while retaining key evolutionary signatures. Together, these results support a mechanism-aware view of protein evolution simulation driven by advanced protein models, in which fitness objectives serve as explicit hypotheses about the molecular constraints that shape evolutionary dynamics.

## Results

To enable sequence simulation explicitly under the constraints provided by different protein models used to evaluate substitution effects, we first implemented a flexible and modular platform named SeqSimFit (**<u>Seq</u>**uence **<u>Sim</u>**ulation under **<u>Fit</u>**ness constraints), shown in Figure 1. In each simulation step, a single amino acid substitution is proposed, the resulting variant is evaluated using a predefined fitness objective derived from one or more protein models, and then the mutation is accepted or rejected by comparing the mutant and current fitness scores under an origin-fixation model (see Materials and Methods). Iterating this procedure generates forward evolutionary trajectories, and the same principle can be applied recursively along a phylogeny by treating each branch as a forward simulation initialized from its ancestral sequence.

**Figure 1.**
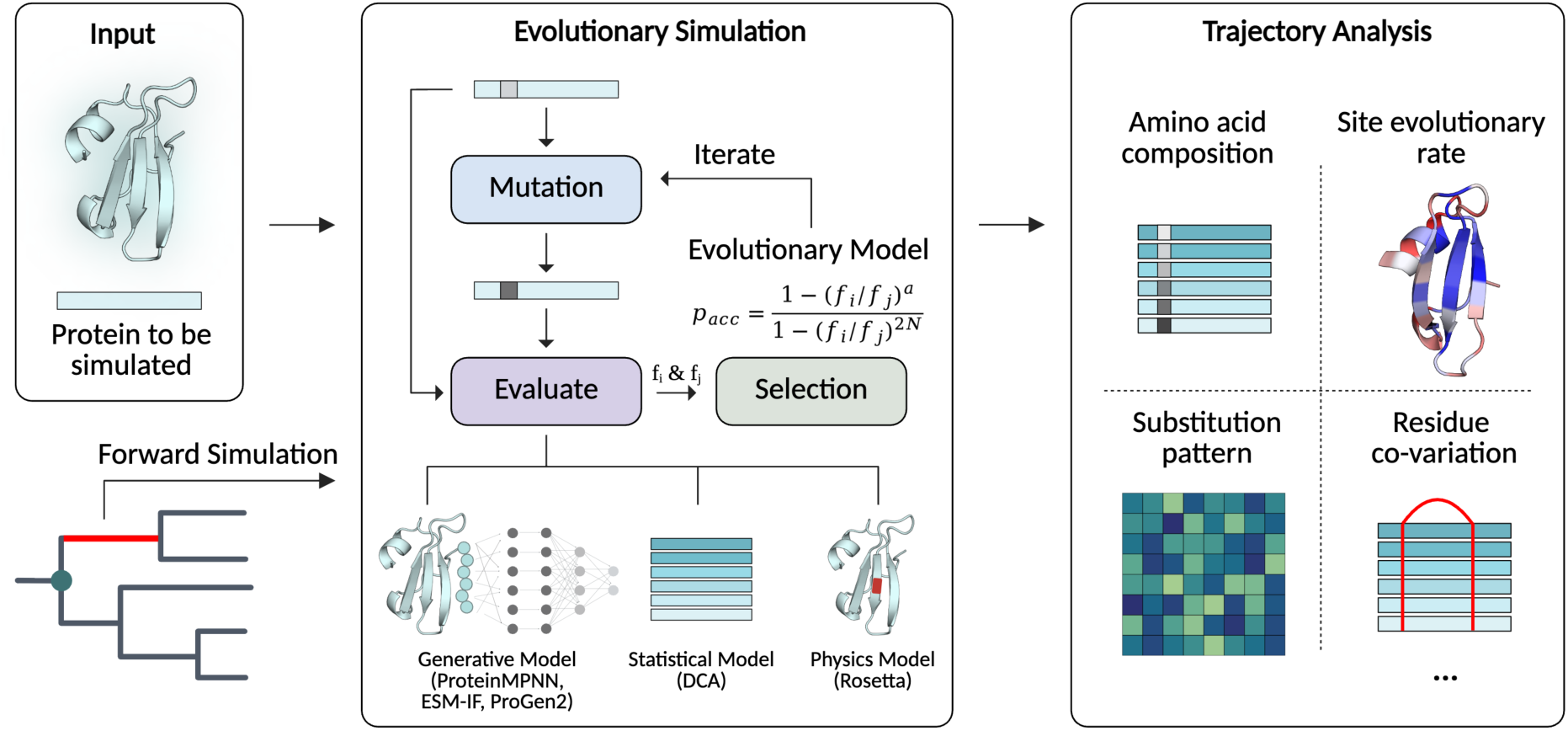
Overview of fitness-guided protein sequence simulation. Starting from an input protein sequence and structure, evolutionary trajectories can be generated either by forward simulation or recursively along a given phylogeny. At each simulation step, a single amino acid substitution is proposed, the mutant sequence is evaluated using a predefined fitness objective, and the mutation is accepted or rejected according to an origin-fixation evolutionary model based on the fitness values before and after mutation. The fitness objective can be derived from different classes of protein models, including generative and inverse-folding models such as ProteinMPNN, ESM-IF, and ProGen2, statistical models such as DCA, and physics-based models such as Rosetta. Simulated trajectories are then analyzed across multiple evolutionary statistics and molecular properties.

For the general evaluation of evolutionary patterns and molecular properties guided by different models (section 1-3 and 6), we compiled a dataset of 100 proteins with lengths between 50 and 200 amino acids, covering diverse structural architectures while maintaining computationally tractable sequence lengths (SI Figure 1). Unless otherwise specified, we generated 50 forward trajectories, each continued until 500 substitutions had been accepted, for each protein under each fitness constraint. These fitness-guided simulations used uniform mutation proposals, with both the sequence position and replacement amino acid sampled uniformly. Five fitness proxies were considered in this work: ProGen2 likelihood (31), ProteinMPNN logits (41), ESM-IF likelihood (42), Rosetta energy score (an approximation of folding ΔG) (28), and DCA statistical energy (29, 43). These proxies represent complementary views of protein constraint: ProGen2 captures protein grammars learned from natural sequences; ProteinMPNN and ESM-IF evaluate sequence–structure compatibility through inverse folding models; Rosetta provides a hybrid knowledge- and physics-based stability estimate; and DCA, fit to natural alignments, serves as an alignment-dependent reference for family-specific evolutionary constraints. As a baseline, we performed empirical substitution simulations using the LG model implemented in Pyvolve under matched trajectory lengths and replicate numbers (19, 22). The oligomeric (section 4) and metamorphic (section 5) protein analyses used separate representative systems, which are introduced in the corresponding sections below and in Materials and Methods.

### 1. Fitness-guided simulations recapitulate multidimensional evolutionary trajectory statistics

To assess whether simulated trajectories capture site-level evolutionary patterns, we first compared their amino acid distributions with natural MSAs using site-wise Kullback–Leibler (KL) divergence, where 0 denotes identical distributions (Figure 2A). As expected, DCA-constrained simulations achieved the lowest average KL divergence (0.51), consistent with their direct dependence on natural MSAs. In contrast, LG-based empirical simulations with Pyvolve showed the highest average divergence from natural MSAs (average KL divergence of 1.14). Among the remaining models, ProteinMPNN generated trajectories closest to natural MSAs (0.96), followed by Rosetta, ProGen2, and ESM-IF (0.99, 1.01, and 1.06, respectively). Pairwise comparisons of per-protein KL divergence scores relative to the corresponding natural MSAs revealed strong concordance among ProteinMPNN, ESM-IF, and Rosetta (Spearman R > 0.5, SI Figure 2). Interestingly, despite showing lower average divergence than the empirical model, ProGen2-guided simulation exhibited the greatest variability across proteins, performing particularly well for some proteins and poorly for others.

**Figure 2.**
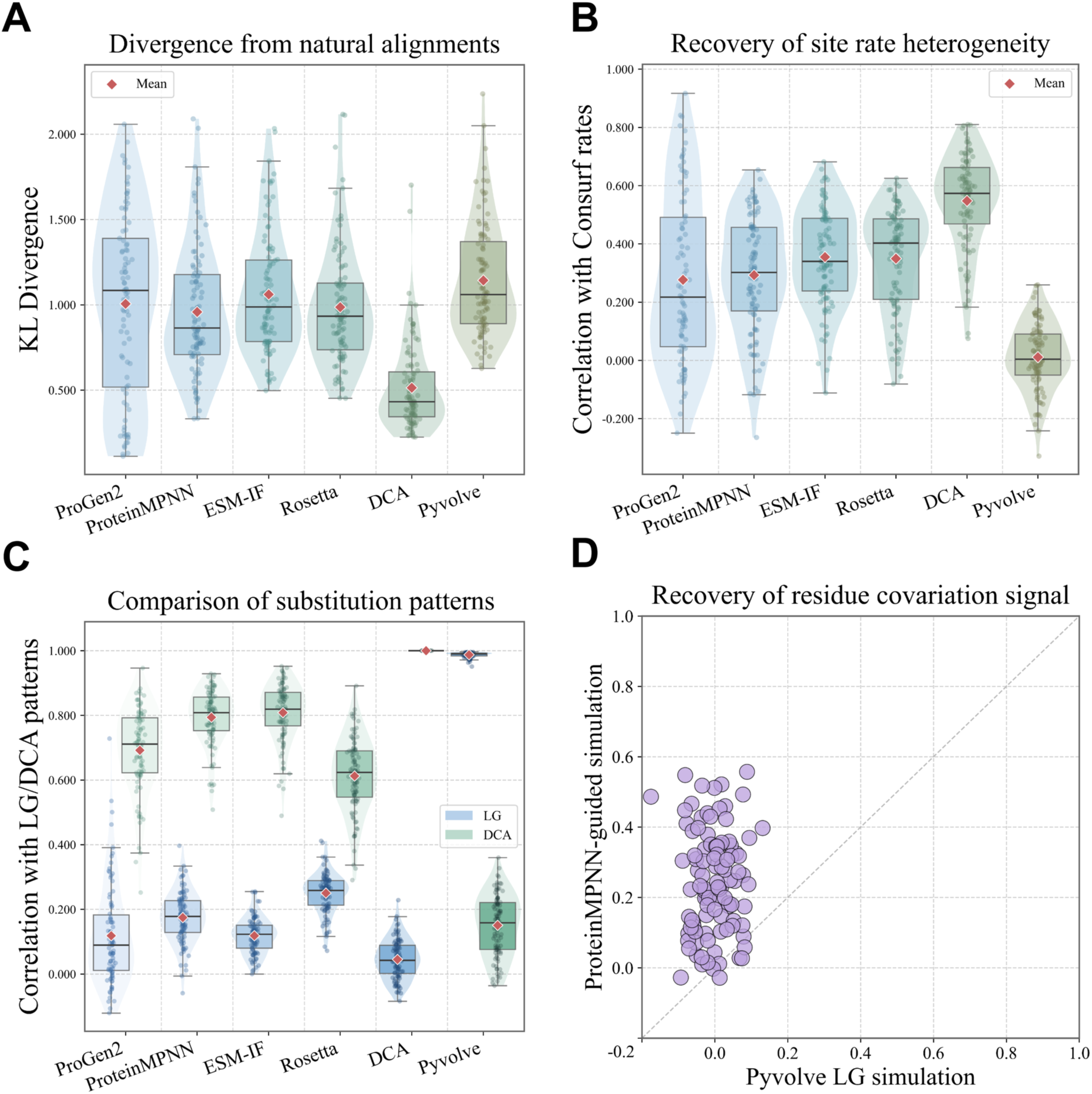
Fitness-guided simulations recover multiple trajectory-level signatures of natural protein evolution. (A) Divergence between simulated trajectories and natural multiple sequence alignments, quantified by site-wise Kullback–Leibler (KL) divergence of amino acid distributions. Lower values indicate closer agreement with natural alignments. (B) Recovery of site-specific evolutionary rate heterogeneity, measured as the Spearman correlation between simulated site substitution counts and ConSurf-derived site rates. Higher values indicate better recovery of natural rate variation. (C) Comparison of substitution patterns inferred from simulated trajectories with either the empirical LG exchangeability matrix or the exchangeability matrix inferred from DCA-guided simulations for the same protein, represented by blue and green boxes, respectively. (D) Recovery of residue– residue co-variation signals. Couplings inferred from ProteinMPNN-guided simulated alignments are compared with those inferred from Pyvolve LG simulations, using natural alignments as the reference. Each point represents one protein. Points above the diagonal indicate stronger recovery under ProteinMPNN-guided simulation. In violin and box plots, individual points represent proteins, red diamonds indicate the mean, boxes show the interquartile range, center lines indicate the median, and whiskers indicate the data range excluding outliers.

We next evaluated whether simulations recover site-specific evolutionary rate heterogeneity (Figure 2B). As a reference, we used rates inferred from natural MSAs and phylogenies by Rate4Site in ConSurf-DB (44, 45), while the simulated trajectory rates were approximated as the number of accepted substitutions at each position. DCA-constrained simulations achieved the highest mean Spearman correlation with natural rates (R = 0.55), again consistent with their use of family-specific MSAs. Empirical Pyvolve simulations without fitness constraints failed to recover natural site-specific rate variation (average R = 0.01), and adding more complex empirical rate models, such as gamma-distributed rates with invariant sites, did not improve performance (SI Figure 3). In contrast, the other fitness-guided simulations partially recovered natural rate variation, with average correlations of 0.28 for ProGen2, 0.30 for ProteinMPNN, 0.35 for ESM-IF, and 0.35 for Rosetta. Pairwise comparisons of per-protein correlations between simulated and ConSurf-derived natural rates further revealed strong agreement among these models (Spearman R > 0.6; SI Figure 2). As an additional sensitivity analysis, we asked whether the strength of the imposed fitness constraint quantitatively affects divergence from natural MSAs and recovery of site-specific evolutionary rate heterogeneity. To this end, using the same ProteinMPNN-guided simulation scheme, we scanned the selection pressure parameter *s* from 0.1 to 1.0 across the same proteins (Materials and Methods). As expected, increasing selection pressure led to a general decrease in divergence from natural alignments and a corresponding increase in the recovery of natural site-specific rates (SI Figure 4). This monotonic trend confirms that the adopted fitness mapping controls the strength of selection in the expected direction, and that stronger constraints produce trajectories closer to natural sequence variation.

We then examined whether fitness-guided simulations produce distinct amino acid substitution preferences (Figure 2C). Substitution matrices were estimated from accepted mutations along simulated trajectories and compared with the global LG exchangeability matrix. As expected, LG-based Pyvolve simulations showed nearly perfect agreement with LG (Spearman R = 0.98). Fitness-guided simulations with a uniform mutation proposal scheme were substantially less correlated with the LG reference, with average correlations of 0.27, 0.26, 0.23, and 0.19 for Rosetta, ProteinMPNN, ProGen2, and ESM-IF, respectively; DCA showed the lowest correlation (Spearman R = 0.12), despite its strong recovery of site-wise evolutionary features (Figure 2A and B). The particularly low DCA-LG correlation is consistent with the difference between global empirical substitution tendencies represented by the LG matrix and the family-specific substitution tendencies derived from DCA models trained on the corresponding protein-family alignments. This motivated us to ask whether fitness-guided simulations could recover the latter. In this comparison, the other fitness-guided methods showed strong agreement with DCA-derived substitution patterns (ESM-IF, 0.81; ProteinMPNN, 0.79; ProGen2, 0.69; Rosetta, 0.61), while Pyvolve remained weakly correlated (R = 0.15). Because substitution patterns inferred from accepted mutations can reflect both the mutation proposal distribution and fitness-based selection, we next assessed the contribution of the proposal scheme. To this end, we repeated ProteinMPNN-guided simulations using LG-based instead of the default uniform mutation proposals. Under this setting, the Spearman correlation with LG increased from 0.26 under uniform proposals to 0.92 (SI Figure 5), indicating that the proposal distribution strongly influences global substitution patterns. However, these trajectories retained key features of fitness-constrained evolution: divergence from natural alignments was maintained or further reduced (average KL divergence, 0.85 vs. 0.96) under the same ProteinMPNN constraint (SI Figure 6), and site-specific rate recovery remained comparable (average R, 0.31 vs. 0.30) to ProteinMPNN simulations with uniform mutation proposals (SI Figure 7). Thus, it is the mutation proposal scheme that largely determines the raw substitution spectrum, whereas the ProteinMPNN-based fitness constraint preserves the improved recovery of site-wise composition and rate heterogeneity.

Finally, we asked whether simulated trajectories encode residue–residue co-variation, a higher-order signal associated with epistatic constraints, which we previously used to assess protein design procedures (46). For each protein, we simulated evolution along phylogenetic trees, collected the leaf sequences to produce alignments, characterized residue couplings, and compared them with those inferred from natural alignments (Materials and Methods). We used phylogeny-based sampling for this analysis because detectable co-variation emerges over extended and diverse evolutionary trajectories (36). Phylogenetic branching provides a practical way to sample such histories, whereas the shorter forward simulations used for the preceding analyses produced substantially weaker co-variation signals. As expected for a site-independent model, LG-based Pyvolve simulations failed to recover natural co-evolution signals, with an average Spearman correlation of 0.01 (Figure 2D). In contrast, fitness-constrained simulation, represented here by ProteinMPNN due to its computational efficiency (SI Table 1), showed substantially stronger agreement with couplings inferred from natural alignments, outperforming Pyvolve for nearly all tested proteins and reaching correlations of up to 0.6 in some cases (Figure 2D).

Together, these analyses show that, compared to empirical matrix-based simulations, fitness-guided simulations improve the recovery of multiple trajectory-level properties of natural evolution, including site-wise amino acid distributions (Figure 2A), site-specific evolutionary rate heterogeneity (Figure 2B), and, in the case of ProteinMPNN, residue-residue co-variation (Figure 2D). This is consistent with previous structure- and fitness-constrained simulation studies showing recovery of site-specific evolutionary patterns and co-variation (33, 47). Importantly, global substitution patterns were largely determined by the mutation proposal scheme (Figure 2C and SI Figure 5), while site-wise composition and rate recovery for ProteinMPNN remained comparable under alternative proposal schemes (SI Figures 6 and 7), indicating that the mutation proposal scheme and fitness-based selection make separable contributions. Finally, different fitness proxies varied in the extent to which they recovered different evolutionary constraints.

### 2. Molecular evaluation reveals consistent drift from wild-type-like properties

Having established that fitness-guided simulations improve trajectory-level realism, we next asked whether the evolved sequences generated along these trajectories also retain favorable molecular properties. Simulations may reproduce natural patterns of site variability or co-variation without necessarily preserving other favorable molecular properties of the corresponding wild-type proteins. We therefore evaluated the evolved sequences using complementary metrics. To compare simulation schemes at a controlled evolutionary distance, for each protein, we collected evolved sequences with 50% identity to the corresponding wild-type protein across all 50 trajectories.

We first assessed sequence plausibility using ESM2 likelihoods expressed relative to those of the corresponding wild-type sequences (Figure 3A). All methods showed average ratios of approximately 1.6-2.1, indicating substantially lower sequence plausibility regardless of the simulation scheme, with ProGen2-guided simulations showing slightly more favorable ESM2 ratios. We then evaluated the structural plausibility of the corresponding ESMFold models (Figure 3B) and found that their pLDDT scores were also reduced relative to the wild type across all simulation schemes, but in this case the differences between schemes were more pronounced. ESMFold models of DCA-guided sequences achieved the highest average pLDDT, followed by models of ProteinMPNN-guided sequences. Interestingly, LG-generated sequences showed higher average predicted structural confidence than those generated under Rosetta, ProGen2, or ESM-IF constraints. The particularly low pLDDT values of ESM-IF-guided sequences were unexpected given its inverse-folding objective, suggesting that iterative optimization under a single sequence– structure compatibility score does not necessarily preserve high predicted structural confidence over long mutational trajectories. Finally, we assessed the energetic compatibility of evolved sequences with the original backbone using DOPE scores (Figure 3C). Differences between simulation schemes were small, and all schemes produced sequences with slightly less favorable DOPE energies than the wild type, with ProteinMPNN showing the most favorable average score.

**Figure 3.**
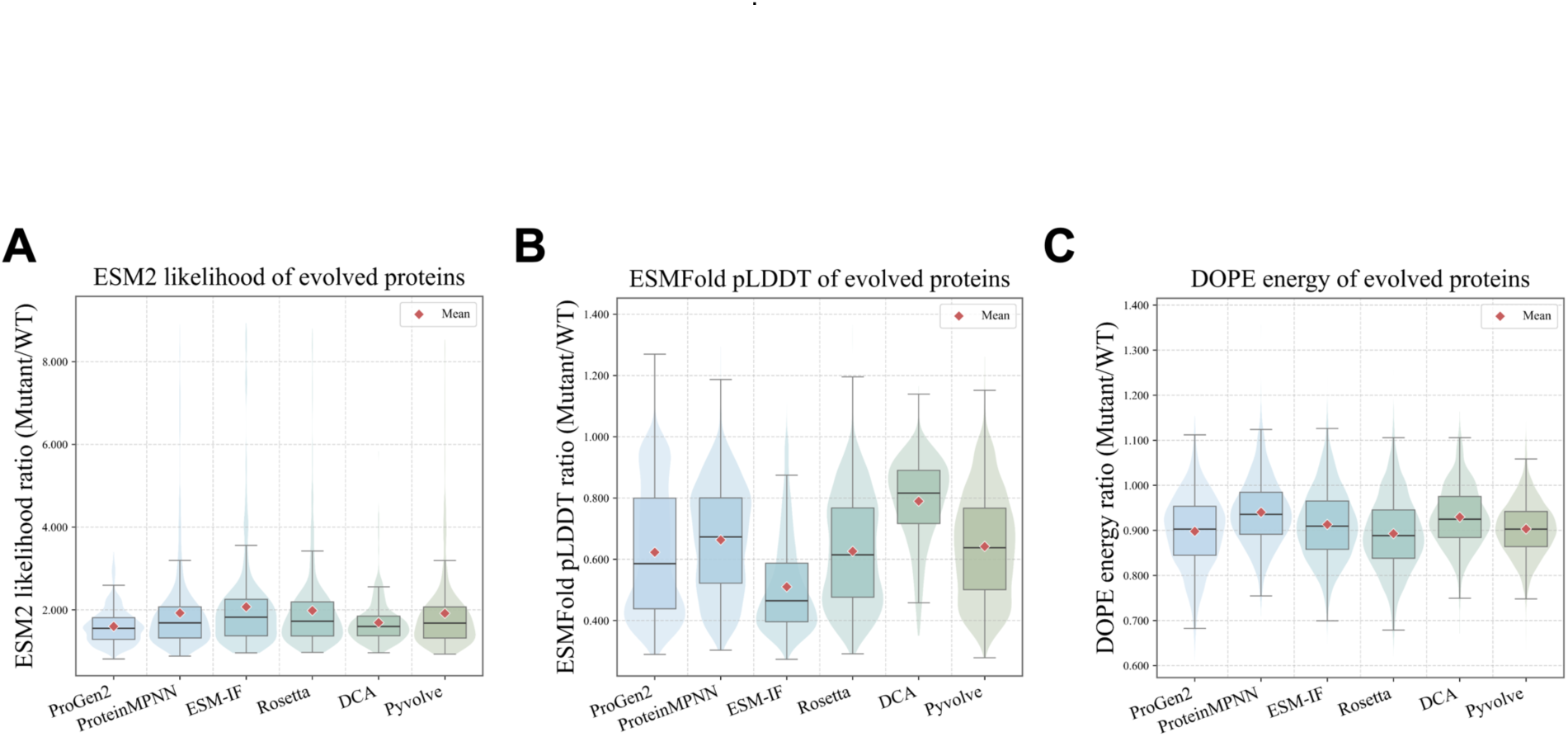
Molecular evaluation reveals objective-dependent trade-offs in evolved sequences. Evolved sequences generated by each simulation strategy were evaluated at 50% sequence identity to their corresponding wild-type proteins. (A) Sequence plausibility measured by ESM2 pseudo-log-likelihood, shown as the ratio between evolved and wild-type sequences. Because ESM2 pseudo-log-likelihood values are negative, ratios closer to 1 indicate scores closer to the wild type, whereas ratios above 1 indicate less favorable scores. (B) Predicted structural confidence measured by ESMFold pLDDT, shown as the evolved/wild-type ratio. Higher ratios indicate better preservation of predicted structural confidence. (C) Energetic compatibility with the wild-type backbone measured by DOPE energy, shown as the evolved/wild-type ratio. Higher ratios indicate more favorable DOPE energies relative to the wild type. In violin and box plots, red diamonds indicate the mean, boxes show the interquartile range, center lines indicate the median, and whiskers indicate the data range excluding outliers.

Together, these molecular evaluations show that, regardless of the simulation scheme used, the resulting sequences are assessed by ESM2, ESMFold, and DOPE as generally less favorable than the corresponding wild-type sequences (Figure 3). Together with the trajectory-level analyses above (Figure 2), this suggests that trajectory realism and molecular properties are non-equivalent: simulations that better recovered evolutionary statistics did not necessarily produce sequences with the best language-model likelihood, predicted structural confidence, or backbone compatibility. However, lower model-derived scores do not necessarily indicate lower molecular quality. Rosetta-designed sequences have previously been shown to receive worse ESM scores than their original sequences despite favorable Rosetta energies (48), while ESMFold confidence only moderately distinguished successful from unsuccessful de novo designs (49).

### 3. Composite fitness objectives balance performance across evolutionary and molecular metrics

The analyses above showed that different fitness proxies produce distinct evolutionary outcomes (Figure 2), while no single proxy preserves wild-type-like molecular properties across the molecular evaluation metrics (Figure 3). Since natural proteins are shaped by multiple constraints simultaneously, these observations motivated us to ask whether complementary protein models could be combined within a programmable fitness objective to produce more balanced simulations. As a proof of concept, we linearly combined scores from three deep learning–based models: ProGen2, ProteinMPNN, and ESM-IF. These models capture distinct signals of protein fitness, with ProGen2 capturing sequence-level plausibility and ProteinMPNN and ESM-IF evaluating sequence–structure compatibility. We applied the same simulation procedure to all seven single-and multi-model scenarios (three single-model, three pairs, and one three-model setup) and evaluated the resulting trajectories using the metrics established above (Figures 2 and 3), including site-wise amino acid composition, site-specific rate recovery, ESM2 likelihood, ESMFold pLDDT, and DOPE energy (Figure 4 and SI Figure 8).

**Figure 4.**
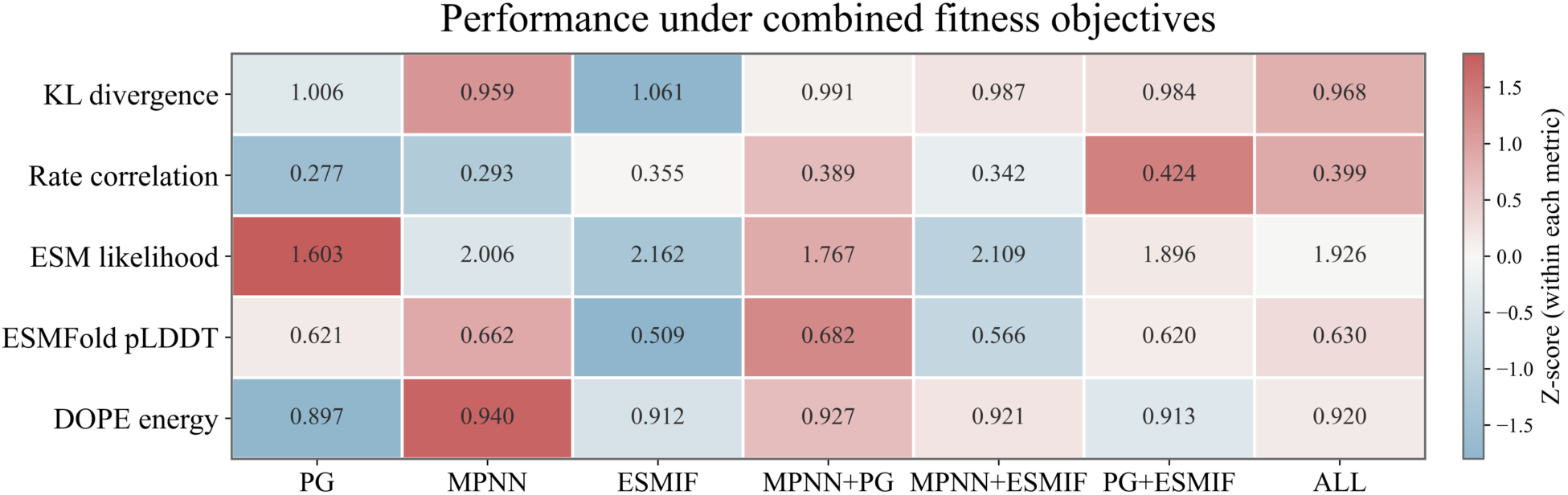
Composite fitness objectives show distinct performance across evaluation metrics. Fitness objectives were constructed from single or linearly combined scores of ProGen2, ProteinMPNN, and ESM-IF, and evaluated using the trajectory-level and molecular metrics established above. Rows show divergence from natural alignments measured by site-wise KL divergence, recovery of site-specific evolutionary rates measured by correlation with ConSurf rates, ESM2 likelihood ratio, ESMFold pLDDT ratio, and DOPE energy ratio of evolved sequences relative to wild-type proteins. Columns represent the seven single- and multi-model objective settings. Cell values show mean raw metric values across the 100-protein benchmark. Cell colors show Z-scores calculated separately within each metric across the seven objective settings, after aligning metric directions so that higher Z-scores consistently indicate better performance (red), whereas lower Z-scores indicate worse performance (blue). Raw distributions are shown in SI Figure 8. PG: ProGen2; MPNN: ProteinMPNN; ESMIF: ESM-IF; ALL: ProGen2+ProteinMPNN+ESM-IF.

Indeed, composite objectives improved several evaluation metrics. For example, although ProGen2 alone showed relatively weak recovery of site-specific rate heterogeneity, combining ProGen2 with ESM-IF increased the average correlation with natural site rates to 0.42, exceeding either individual model (0.28 and 0.35, respectively). Also, a slight improvement in predicted structural confidence was observed for the ProGen2-ProteinMPNN combination, which achieved an average pLDDT ratio of 0.682 compared with 0.621 for ProGen2 and 0.662 for ProteinMPNN alone. These improvements indicate that in some cases composite objectives can integrate useful signals from different modeling paradigms, allowing sequence-level (ProGen2) and structure-level (ESM-IF, ProteinMPNN) constraints to jointly shape simulated evolution. However, multi-model constraints did not uniformly outperform the best single-model objectives. For sequence-related metrics such as alignment divergence and ESM2 likelihood, the single-model ProteinMPNN and ProGen2 constraints, respectively, remained the best-performing scenarios. Similarly, ProteinMPNN alone also achieved the lowest average DOPE energy. Thus, model combination should not be interpreted as a simple route to universal improvement. Its main effect is instead to redistribute performance across different evaluation metrics.

A clear example of this balancing effect was the ProGen2-ProteinMPNN simulation. ProGen2 alone produced the best ESM2 likelihood but the worst DOPE energy, whereas ProteinMPNN alone achieved the best DOPE energy but substantially worse ESM2 likelihood. Both also performed poorly in site-specific rate recovery. Their combination largely retained the strengths of both models, preserving high ESM2 likelihood and favorable DOPE energy while improving rate recovery and slightly improving pLDDT relative to either single model. Thus, combining complementary fitness proxies can mitigate the specific weaknesses of individual models without fully sacrificing their strengths.

Together, these results show that the fitness objective is not fixed, but programmable. Rather than selecting a single protein model as the universally best proxy for evolution, different objectives can be combined to impose different combinations of selective constraints. This provides a practical bridge from model comparison toward mechanism-aware simulations in which the fitness objective is constructed to represent specific biological hypotheses.

### 4. Encoding oligomeric context reveals interface-specific selective pressure

We next asked whether such programmable fitness objectives could encode explicit biological context, rather than only combine different model scores. Many proteins function as oligomeric assemblies, where oligomerization interfaces impose structural and evolutionary constraints that may be missed by simulations based on isolated monomer structures. Because ProteinMPNN can evaluate sequence compatibility in multi-chain structural contexts, it allows oligomeric assembly to be incorporated into fitness-guided evolution. Thus, ProteinMPNN can be used to construct programmable fitness objectives in which the relative contributions of monomeric and oligomeric structural contexts are explicitly varied.

To test this idea, we focused on a set of eleven well-characterized homodimeric proteins with biologically relevant dimer architectures (Figure 5A). Using ProteinMPNN as the fitness evaluator, we simulated evolutionary trajectories under structural constraints ranging from the isolated monomeric structure extracted from the corresponding dimer to the full dimeric assembly. Specifically, this continuum was generated by varying the relative weights of ProteinMPNN scores calculated from the monomeric and dimeric structural contexts. We then evaluated the resulting trajectories using the metrics introduced above, focusing on the recovery of site-wise amino acid composition and site-specific evolutionary rates. Because these measures exhibited a roughly monotonic trend across the weighting continuum in many cases (SI Figure 9), we present only the two extreme cases (Figure 5B-D), i.e., purely monomeric and purely dimeric constraints, for clarity.

**Figure 5.**
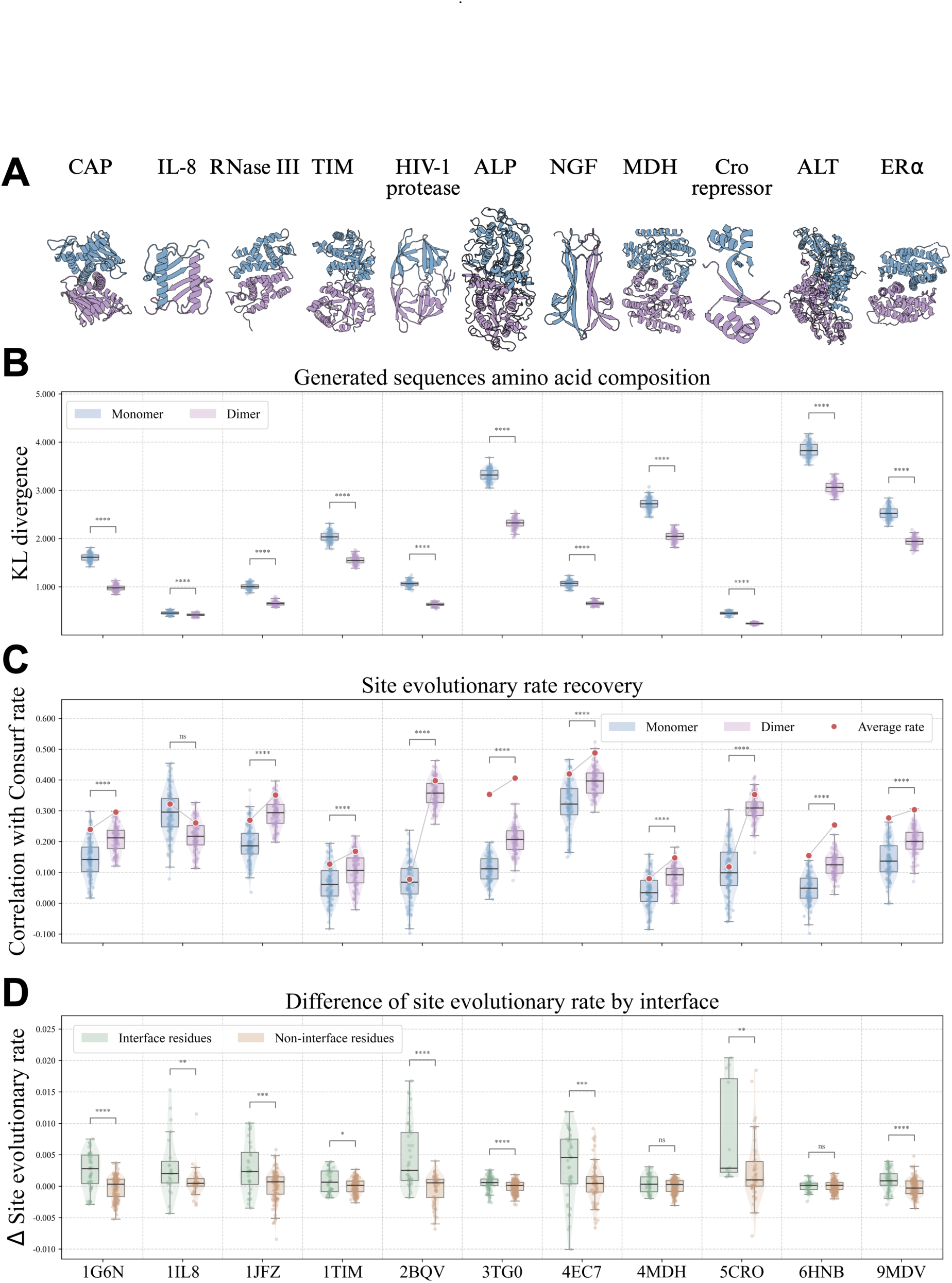
Encoding oligomeric context reveals interface-specific selective pressure in homodimeric proteins. (A) Representative biologically relevant homodimeric structures used for monomer- and dimer-constrained simulations. Chains are shown in different colors. (B) Divergence between simulated trajectories and natural alignments, measured by site-wise KL divergence of amino acid distributions. For each protein, ProteinMPNN-guided simulations were performed using either the isolated monomeric structure or the full dimeric assembly as the structural context. Lower KL divergence indicates closer agreement with natural sequence variation. (C) Recovery of site-specific evolutionary rate heterogeneity, measured as the correlation between simulated site substitution counts and ConSurf-derived site rates. Red points indicate the correlation computed from the average simulated site-rate profile across trajectories. (D) Interface-specific changes in simulated evolutionary rates after incorporating dimeric context. Residues were classified as interface or non-interface sites, and the change in site evolutionary rate between monomer- and dimer-constrained simulations was compared. Positive values indicate stronger rate reduction under dimer-aware simulation. Significance was assessed using the Mann–Whitney U test; ns, not significant; *p < 0.05; **p < 0.01; ***p < 0.001; ****p < 0.0001. Box plots show the interquartile range and median, with individual points representing simulated trajectories or residues as appropriate.

Incorporating dimeric structural context consistently improved agreement between simulated and natural sequence variation. Across all tested proteins, dimer-aware simulations showed significantly lower divergence from natural alignments than monomer-based simulations (Figure 5B, p < 1e-4, Mann–Whitney U test). Consistently, the recovery of natural site-specific evolutionary rates was also markedly improved for all proteins except interleukin-8 (Figure 5C, p < 1e-4, Mann–Whitney U test). In HIV-1 protease, for example, the correlation with ConSurf-derived rates increased from 0.05 under monomer-based simulation to 0.41 when dimer information was included, consistent with the fact that it requires the dimeric state to form a functional active site (50). In contrast, IL-8 exists in a dynamic monomer–dimer equilibrium and its monomer is highly relevant for receptor activation, while the dimer contributes to glycosaminoglycan binding and neutrophil recruitment (51–53). Consistent with the functional relevance of both oligomeric states, IL-8 was the only protein for which the fully dimeric constraint reduced site-rate recovery relative to the monomeric context, while a mixed monomer-dimer weighting yielded the highest rate recovery (Figure 5C; SI Figure 9B).

We then asked whether this improvement reflected biologically interpretable changes in the simulated trajectories. Because interface residues are expected to experience stronger constraints than non-interface residues, we compared site-specific rate changes between dimer- and monomer-based simulations after classifying residues by interface involvement. In 9 out of 11 proteins, incorporating dimer constraints led to a significantly larger decrease in evolutionary rates at interface residues than at non-interface residues (Figure 5D, p < 1e-2, Mann–Whitney U test). Thus, the effect of adding dimeric structural context was not merely global, but was preferentially localized to residues involved in the dimer interface.

To test whether the improvement under dimeric constraints (Figure 5) depended on the biologically correct interface, rather than simply on the addition of any oligomeric context, we generated three alternative dimer decoys for each of 11 proteins using HDOCK (SI Figure 10, Materials and Methods) (54). The selected decoys were structurally plausible but clearly distinct from the corresponding experimental dimeric structures, and were subjected to the same simulation and analysis procedure used in Figure 5. Interestingly, most decoy dimers produced lower KL divergence from natural alignments than the corresponding monomer-based simulations, although the experimentally determined dimers produced the closest agreement in most cases (SI Figure 11), initially suggesting that the observed effect cannot be solely attributed to the introduction of any dimer partner. However, examination of sequence identity relative to the starting wild-type sequence revealed a similar ordering, with monomers generally exploring sequence space most extensively, followed by decoys, and native dimers being the most constrained (SI Figure 12). Notably, IL-8 and TIM showed the same deviation from this typical ordering in both analyses, i.e., some decoys that restricted sequence exploration as strongly as (or more strongly than) the native dimer also showed comparable or lower KL divergence. This suggests that the introduction of an additional intermolecular interface can generically restrict sequence exploration around the wild-type sequence, which may in turn partially account for the lower KL divergence without necessarily reflecting the biological correctness of the interface.

For site-specific evolutionary rate recovery, we also saw a clear pattern suggesting that simulations using the native dimer more accurately recover natural site-rate profiles: decoy-based simulations did not systematically outperform the monomer-based simulations and consistently showed weaker agreement with natural profiles than simulations using the native dimer structures (SI Figure 11). Importantly, unlike the KL-divergence result, this pattern does not appear to be explained simply by a generic restriction of sequence exploration because decoys that restricted sequence exploration as strongly as the native dimer still showed poorer recovery of natural site-rate profiles. This is consistent with site-rate recovery depending not only on the overall strength of constraint but also on where that constraint is imposed along the sequence, making it more sensitive to whether the modeled interface corresponds to the biologically relevant one. We note, however, that decoy dimers cannot be perfectly matched to the native assembly in properties such as interface size and contact number, and may also partially reuse native interface residues. A fully controlled comparison would therefore require more systematic decoy design in future work.

Taken together, by changing the structural context provided to the same fitness model, the simulations captured a key mechanistic feature of protein evolution, whereby residues participating in biologically relevant interfaces are selectively constrained and therefore evolve more slowly. This supports the broader idea that programmable fitness objectives can be used not only to improve agreement with natural sequence variation, but also to test how specific biological constraints shape evolutionary trajectories.

### 5. Multi-conformation fitness objectives capture evolutionary constraints in metamorphic proteins

Motivated by the results from the dimer-aware simulations, we then asked whether this idea could be extended to proteins whose evolutionary constraints may not be fully represented by a single native conformation. Such metamorphic proteins reversibly switch between distinct folds under physiological conditions and therefore challenge the conventional one-sequence–one-structure view (55). They provide a stringent test case: if multiple conformations impose evolutionarily relevant constraints, simulations incorporating both states should better reproduce natural sequence variation than simulations constrained by either state alone.

We first examined XCL1, a well-characterized metamorphic chemokine that interconverts between an α+β chemokine fold (1J8I) (56) and an all-β dimeric state (2JP1) (57) (Figure 6A). We defined the fitness objective as a weighted linear combination of ProteinMPNN-derived fitness terms computed for the two conformations in their biologically relevant oligomeric contexts and simulated evolution across weighting schemes ranging from the all-β-fold-only constraint (*w_A_* = 0.0, *w_B_* = 1.0) to the chemokine-fold-only constraint (*w_A_* = 1.0, *w_B_* = 0.0) with A and B denoting the chemokine and all-β states, respectively. The resulting trajectories were again evaluated using site-specific amino acid distributions and site-specific rate heterogeneity.

**Figure 6.**
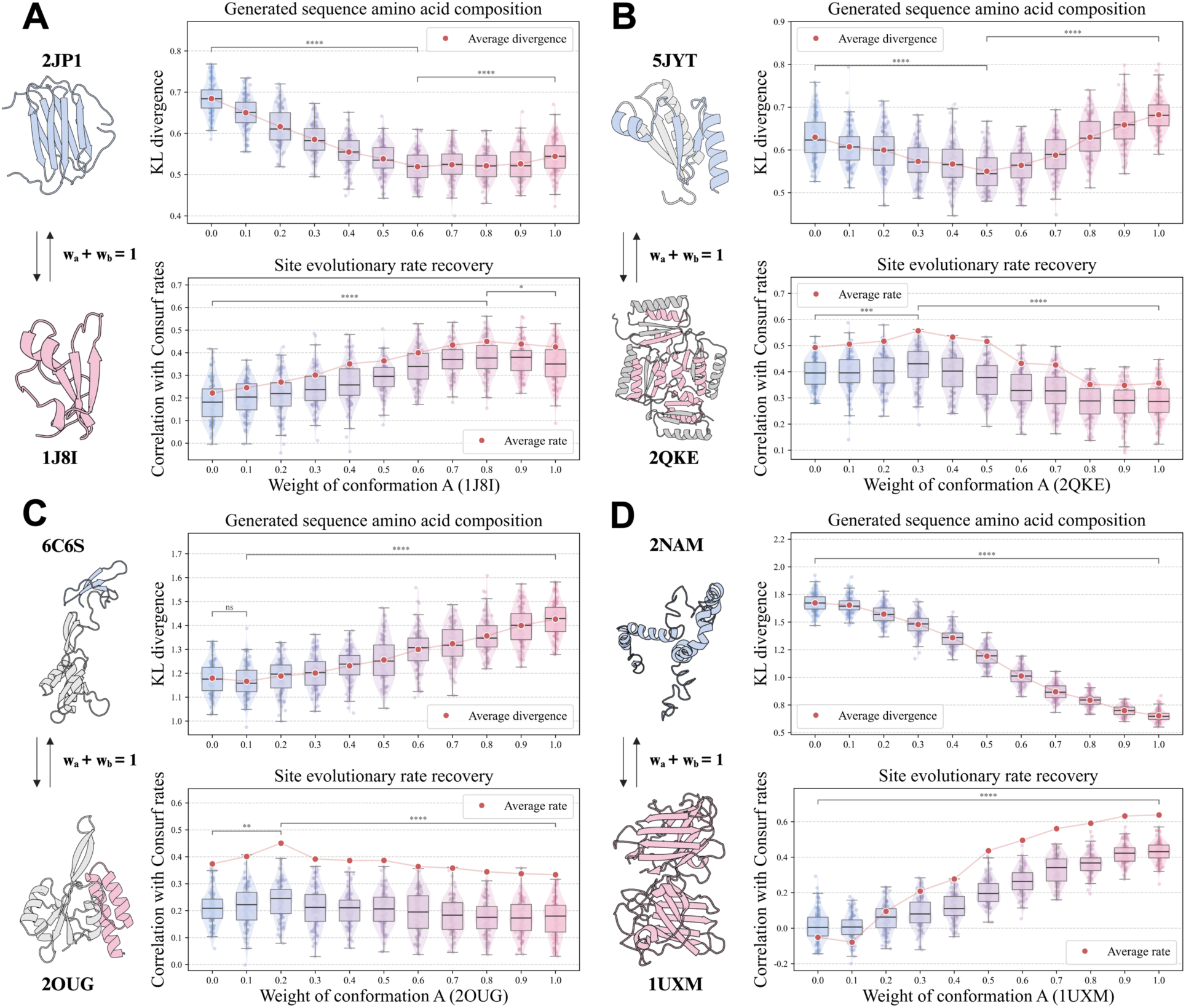
Multi-conformation fitness objectives capture evolutionary constraints in metamorphic proteins. For each protein, two experimentally characterized conformational states were used to define a weighted ProteinMPNN-based fitness objective, and simulations were performed across different relative weights assigned to the two conformations (*w_a_* + *w_b_* = 1). (A) XCL1, with conformation A corresponding to the α+β chemokine fold (PDB: 1J8I). (B) KaiB, with conformation A corresponding to the ground-state C-terminal fold (PDB: 2QKE). (C) RfaH, with conformation A corresponding to the α-helical C-terminal fold (PDB: 2OUG). (D) SOD1, with conformation A corresponding to the native β-barrel fold (PDB: 1UXM). For each system, the upper plot shows divergence from natural alignments measured by site-wise KL divergence of amino acid distributions, and the lower plot shows recovery of site-specific evolutionary rate heterogeneity measured by correlation with ConSurf-derived rates. Red points indicate aggregate divergence or rate-correlation values computed after pooling sequences from all 100 simulated trajectories. Statistical comparisons were performed using the Mann–Whitney U test; ns, not significant; *p < 0.05; **p < 0.01; ***p < 0.001; ****p < 0.0001. In violin and box plots, individual points represent trajectories, boxes show the interquartile range, center lines indicate the median, and whiskers indicate the data range excluding outliers.

The simulations showed a strong dependence on the relative weighting of the two conformations (Figure 6A). Simulations constrained only by the all-β fold (*w_A_* = 0.0, *w_B_* = 1.0) produced the largest divergence from natural XCL1 alignments, whereas increasing the weight of the chemokine fold (*w_A_*) progressively improved agreement with natural sequence variation. The best match was achieved under a mixed objective, with the chemokine fold receiving a dominant but not exclusive weight (*w_A_* = 0.8, *w_B_* = 0.2), which produced significantly lower divergence than either single-conformation extreme (p < 1e-4, Mann–Whitney U test). Site-specific rate recovery showed the same optimum, with the highest correlation to ConSurf-derived rates also occurring at *w_A_* = 0.8 and *w_B_* = 0.2. These results suggest that XCL1 evolution is better modeled by an asymmetric combination of conformational constraints than by either fold alone, consistent with XCL1 having evolved from a monomorphic chemokine-fold ancestor before acquiring metamorphism (58).

Encouraged by these findings, we further extended this analysis to two additional well-studied fold-switching proteins, KaiB and RfaH (Figure 6B and C, respectively) (55). KaiB, a core component of the cyanobacterial circadian oscillator, adopts two interconvertible conformations in its C-terminal region: a ground-state fold with βααβ topology (2QKE) (59) and a fold-switched state with αββα topology (5JYT) (60), corresponding to tetrameric and monomeric contexts, respectively (Figure 6B). As in XCL1, the KL divergence between simulated and natural KaiB sequences exhibited a valley-shaped dependence on the weighting scheme, with the closest agreement at *w_A_* = 0.5 and *w_B_* = 0.5. However, the site-specific rate recovery peaked at *w_A_* = 0.3 and *w_B_* = 0.7, indicating that the rate-recovery optimum favored the fold-switched αββα topology (60). RfaH belongs to the NusG family of transcription elongation factors, with a C-terminal domain capable of switching between an α-helical hairpin fold in the unbound autoinhibited state (2OUG) (61) and a β-roll fold in the activated transcription elongation complex (6C6S) (62). In our simulations, it showed a less pronounced but still informative trend (Figure 6C): incorporating both conformations produced the lowest sequence divergence at *w_A_* = 0.1 and *w_B_* = 0.9, representing a predominantly β-roll-state constraint, with a significant improvement over the α-hairpin-only condition (*w_A_* = 1.0, *w_B_* = 0.0) and only a marginal, statistically non-significant improvement over the β-roll-only condition (*w_A_* = 0.0, *w_B_* = 1.0). Site-specific rate recovery showed a similar but distinct optimum at (*w_A_* = 0.2, *w_B_* = 0.8), significantly outperforming both single-state simulations.

As an additional comparison, we examined human superoxide dismutase (SOD1) (Figure 6D). Under physiological conditions, SOD1 adopts a stable native β-barrel fold (63), whereas an alternative disordered, helix-rich state has been associated with non-native and cytotoxic behavior (64). In contrast to three metamorphic proteins described above, simulations including both SOD1 states did not show an optimal mixed-conformation regime. Instead, sequence divergence decreased and rate recovery improved monotonically as the weight on the native β-barrel state increased. This suggests that adding a non-native conformational state does not by itself generate signatures consistent with fold-switching. We next asked whether they could emerge when the alternative structure was instead a computationally generated, yet structurally plausible, same-fold backbone variant. To this end, for XCL1, KaiB, and RfaH, we paired one experimentally determined conformation with independently generated structural decoys using Rosetta (SI Figure 13, Materials and Methods) and repeated the weighting analysis. Compared with the experimentally characterized fold-switching pairs, these simulations showed substantially less pronounced valley-shaped behavior in sequence divergence and correspondingly weaker peaks in site-specific rate recovery (SI Figure 14). Together, these controls indicate that the observed improvement under multi-conformation objectives cannot be explained simply by adding a second structure or averaging over alternative backbone snapshots, but instead depends on the second conformation representing a biologically meaningful evolutionary constraint.

Collectively, these results demonstrate that programmable fitness objectives can model evolutionary constraints arising from multiple conformational states. Unlike DCA-based simulations, which infer constraints from natural alignments, multi-conformation structure-aware simulations here allow specific structural hypotheses to be encoded directly into the evolutionary process. The optimal weighting schemes suggest that both conformations can impose selective constraints during evolution and may provide a model-dependent readout of the relative contribution of each conformational constraint to reproducing natural sequence variation.

### 6. Differentiable structural constraints improve molecular plausibility while retaining evolutionary signatures

The preceding sections showed that programmable fitness objectives can encode specific structural constraints into simulations of sequence evolution. However, the evaluation of evolved sequences also revealed an important limitation: trajectories that reproduce natural evolutionary statistics can still generate sequences with compromised molecular properties, depending on the fitness proxy used (Figure 3). We therefore asked whether structure-preserving constraints could be incorporated more directly into evolutionary simulation to improve the structural plausibility of evolved sequences.

A natural candidate is AlphaFold2 (65), whose confidence metrics, such as pLDDT and PAE, have been widely used to guide protein design (66, 67). Directly embedding structure prediction into a stepwise mutation–selection procedure is computationally expensive, because each proposed mutation would require repeated structure-prediction evaluation. We therefore adopted an alternative strategy inspired by recent protein design approaches: backpropagation through AlphaFold2 (AF-back) to optimize sequence profiles at the PSSM level (66) (Figure 7A). The procedure starts with a PSSM initialized from the starting sequence and a target backbone that remains fixed throughout the optimization. The sequence profile is used as input to AlphaFold2 to predict a structure, which is evaluated using a composite loss function comprising an RMSD-to-target term, pLDDT, and a contact term. The loss is then backpropagated through AlphaFold2 to obtain gradients with respect to the sequence profile, which are used to optimize the PSSM. This cycle is repeated multiple times, yielding an optimization trajectory. Although AF-back approaches have been used for design, whether such trajectories retain plausible evolutionary signatures remains unclear. To this end, we applied AF-back to the 100-protein dataset and evaluated the resulting trajectories using the same evolutionary and molecular metrics described above.

**Figure 7.**
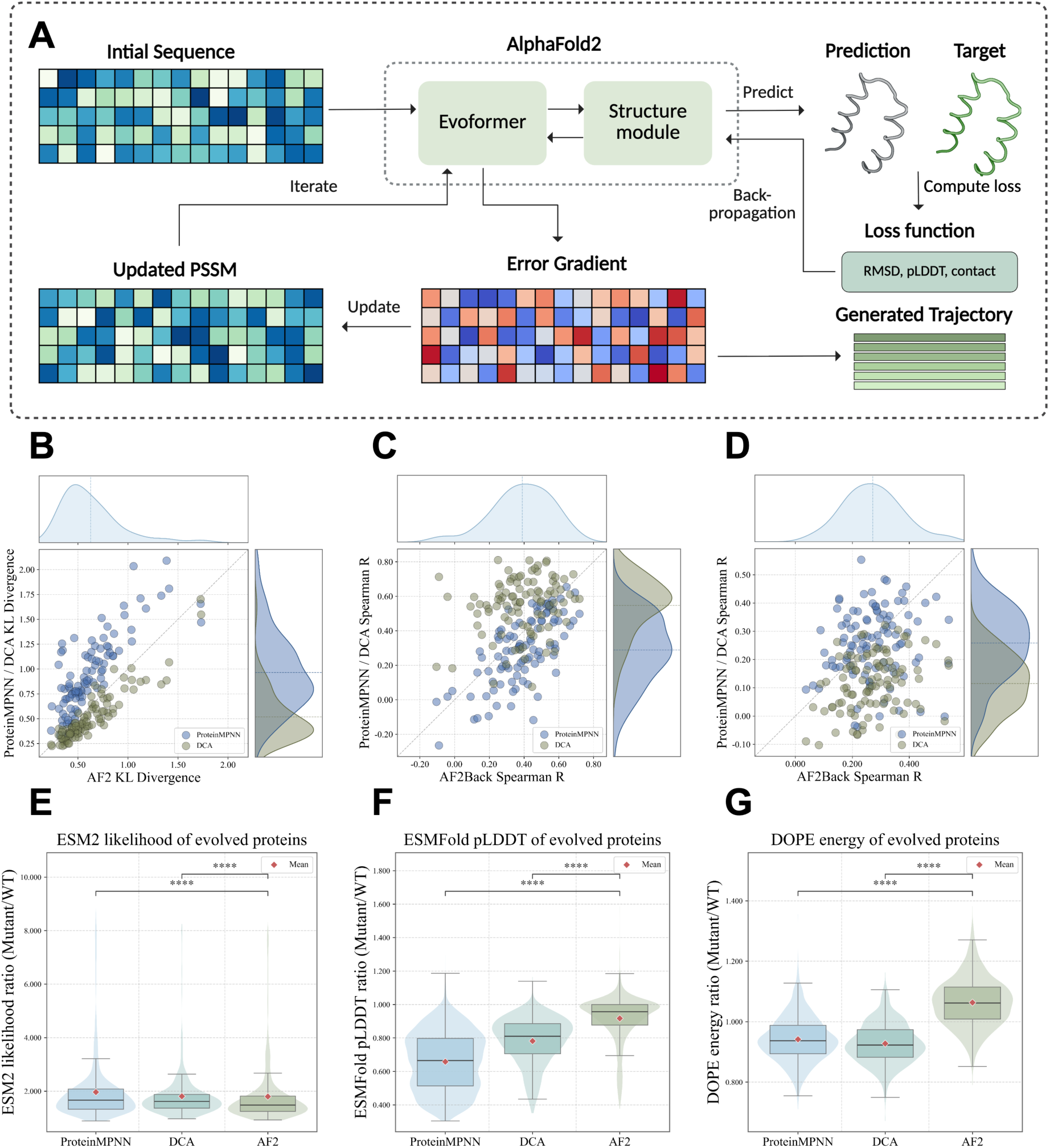
AlphaFold backpropagation provides a differentiable extension of structure-constrained evolutionary simulation. (A) Schematic of the AlphaFold-backpropagation (AF-back) procedure. Starting from an initial sequence profile, AlphaFold2 predicts a structure, which is compared with the target structure using a customized loss function including RMSD, pLDDT, and contact terms. Gradients are backpropagated through the network to update the sequence profile, generating an optimization trajectory that can be analyzed as a structure-constrained sequence trajectory. (B–D) Comparison of AF-back trajectories with ProteinMPNN- and DCA-guided simulations across trajectory-level evolutionary statistics. Each point represents one protein, with AF-back performance shown on the x-axis and the corresponding ProteinMPNN or DCA performance shown on the y-axis. The diagonal indicates equal performance. Marginal density plots summarize the distributions across proteins. (B) Divergence from natural alignments measured by site-wise KL divergence. (C) Recovery of site-specific evolutionary rate heterogeneity measured by Spearman correlation with ConSurf-derived rates. (D) Comparison of substitution patterns inferred from simulated trajectories with the empirical LG exchangeability matrix. (E–G) Molecular properties of evolved sequences generated at 50% sequence identity to the wild type. (E) ESM2 likelihood ratio, (F) ESMFold pLDDT ratio, and (G) DOPE energy ratio of evolved sequences relative to wild-type proteins. Because ESM2 likelihood values are negative, higher evolved/wild-type ratios indicate less favorable likelihoods relative to the wild type. Higher ratios indicate more favorable DOPE energies relative to the wild type. Statistical significance was assessed using the Mann–Whitney U test; ns, not significant; *p < 0.05; **p < 0.01; ***p < 0.001; ****p < 0.0001. In violin and box plots, individual points represent proteins, red diamonds indicate means, boxes show the interquartile range, center lines indicate medians, and whiskers indicate the data range excluding outliers.

We first examined whether AF-back trajectories reproduced trajectory-level statistics of natural evolution by comparing AF-back, ProteinMPNN-, and DCA-guided simulations against the corresponding natural data. AF-back achieved an average KL divergence of 0.63 relative to natural alignments, outperforming ProteinMPNN (0.96) and approaching DCA (0.51) (Figures 7B and 2A). It also exhibited a moderate recovery of site-specific evolutionary rate heterogeneity, reaching an average Spearman correlation of 0.39, again surpassing ProteinMPNN (0.30) and second only to DCA (0.55; Figures 7C and 2B). Given the long-standing hypothesis that structural contacts are key determinants of site variability, we repeated the simulation without the contact term in the loss function. Removing this term reduced rate recovery from 0.39 to 0.27 (SI Figure 15), indicating that the contact term contributes to the recovery of natural site variability in AF-back trajectories.

AF-back trajectories also showed substitution patterns similar to those of other fitness-guided simulations, deviating substantially from the global LG model, with an average correlation of 0.27 (Figures 7D and 2C). We further generated phylogenetically structured alignments (Figure 1) using AF-back and inferred residue–residue couplings from the resulting leaf sequences. AF-back-derived alignments showed slightly stronger co-variation signals than LG-generated alignments (average Spearman R, 0.07 vs 0.00, respectively; SI Figure 16), although considerably weaker than ProteinMPNN-guided alignments (average Spearman R, 0.24; SI Figure 17). We hypothesize that epistatic signals captured by AlphaFold2 are reflected in simulated sequence variation, although to a limited extent, presumably due to the conversion of optimized PSSM sequence profiles to discrete sequences for downstream analysis.

We next evaluated sequences generated at 50% identity to the wild-type as before. In contrast to several discrete fitness-guided simulations, AF-back produced evolved sequences with comparatively favorable scores across all three evaluation dimensions. AF-back-generated sequences achieved better ESM2 likelihood scores than all previously tested simulation strategies except for ProGen2-guided simulations, including DCA-guided simulations (Figures 7E and 3A). They also maintained the highest ESMFold pLDDT values, approaching those of native proteins and exceeding the best discrete mutation–selection simulations, which were DCA-guided (Figures 7F and 3B). Although this improvement is expected because confidence-related terms were included in the optimization loss, it demonstrates that differentiable structural constraints can reduce the loss of predicted structural confidence observed under some single-proxy discrete simulations. Finally, AF-back-generated sequences achieved highly favorable DOPE energies when modeled using the wild-type structure as a template (Figures 7G and 3C), outperforming all other methods and even producing lower average energies than the native sequences themselves, with an average evolved/wild-type ratio of 1.06.

Together, these results show that AlphaFold2 backpropagation provides a complementary route for structure-aware sequence simulation. By borrowing differentiable optimization strategies from protein design, AF-back incorporates structural constraints more directly than the discrete fitness proxies tested above, improving molecular evaluation scores while retaining several evolutionary signatures. At the same time, gradient-based profile optimization is not equivalent to a standard mutation–selection process, and should therefore be interpreted as a differentiable extension rather than a replacement for discrete evolutionary simulation. More broadly, this analysis illustrates a productive connection between protein design and evolution: design methods provide increasingly powerful ways to define molecular constraints, while evolutionary simulation asks whether those constraints generate natural-like patterns of sequence variation.

## Discussion

In this study, we used flexible and programmable fitness objectives driven by diverse protein models to simulate protein evolution under explicit constraints. The central idea is that a fitness objective is not merely a score used to accept or reject mutations, but a hypothesis about what selection preserves. By changing the objective, we can ask how different assumptions about sequence naturalness, structural compatibility, physical stability, oligomeric assembly, or conformational switching shape simulated evolutionary trajectories. This perspective extends protein sequence simulation beyond generic substitution rules, making it possible to test whether proposed molecular constraints are sufficient to reproduce the statistical signatures observed in natural evolution.

Our benchmarks across diverse proteins first established a diagnostic basis for assessing the applicability of fitness objectives derived from different classes of protein models in evolutionary trajectory simulations (Figure 2). Such fitness-guided simulations recovered several trajectory-level features that were otherwise poorly captured by the empirical LG substitution model alone, while different fitness objectives recovered different evolutionary signatures to different extents. Specifically, DCA-based simulations performed strongly because they directly encode family-specific constraints learned from natural alignments, whereas structure-based (ProteinMPNN and ESM-IF), physics- and knowledge-based (Rosetta), and language-model-based (ProGen2) objectives captured different aspects of protein constraint without requiring a family-specific MSA for the target protein. Molecular evaluation further showed that natural-like trajectory signatures do not necessarily translate into favorable sequence likelihood, structural confidence, or backbone compatibility, as assessed by ESM2 likelihood, ESMFold pLDDT, and DOPE energy, respectively (Figure 3). Thus, rather than identifying a universally best fitness proxy, these analyses show that each objective represents a particular proxy for, or hypothesis about, the constraints imposed by selection. Programmability becomes useful in this context because it enables the construction of composite objectives that can combine complementary constraints within a single fitness function (Figure 4). Therefore, this strategy serves less as an optimization exercise than as a bridge toward simulations designed around specific evolutionary hypotheses.

This programmable fitness view became most informative in biological contexts where the relevant constraints are not fully described by a single monomeric fold. In homodimeric proteins, incorporating the oligomeric structure into the same ProteinMPNN-based fitness model improved agreement with natural sequence variation and, more importantly, produced interface-localized changes in evolutionary rates (Figure 5). This is further highlighted by simulations of metamorphic proteins in which a single sequence must remain compatible with multiple native conformations with each conformation represented in its biologically relevant oligomeric context (Figure 6). Multi-conformation fitness objectives allowed us to test whether different structural states contribute to the evolutionary constraints observed in natural sequences. Across the tested fold-switching proteins, mixed-conformation objectives often better reproduced natural sequence variation than objectives constrained by either conformation alone. Because experimentally characterized metamorphic proteins with well-resolved alternative structures remain limited in number, our analysis was restricted to a small set of representative systems. As more candidate metamorphic proteins are identified, this strategy could provide an evolutionary criterion for evaluating the proposed conformational states. In contrast to recent work using inverse folding models to engineer preferences between alternative folds (68), the proposed approach asks whether those folds leave detectable signatures in natural sequence evolution. Together, the dimer and the metamorphic protein analyses show that encoding higher-level structural context in the fitness objective can produce biologically interpretable changes in simulated evolutionary dynamics. This, in turn, suggests that the proposed simulation could be extended to more complex assemblies, such as heterodimers and transient protein– protein interactions, including those coupled to conformational changes. These simulations could also be used to examine the emergence of protein–protein interactions, complementing recent advances in library-based directed evolution (69).

We further showed that AlphaFold2-backpropagation can provide a more direct and effective way of introducing structure-preserving constraints into simulations (Figure 7). However, because simultaneous optimization of the full sequence profile is not equivalent to the stepwise mutation-selection procedure, these trajectories should be interpreted as a complementary extension rather than as direct simulation of the standard evolutionary process. Beyond the structure-preservation objective explored in this work, the AF-back objective could in principle be easily adapted to encode more specific and mechanism-aware molecular constraints. For example, the success of AlphaFold-backpropagation-based binder design pipelines such as BindCraft suggests the possibility of simulating protein evolution under explicit interaction or binding-related constraints (70). Echave, meanwhile, demonstrated that accounting for both stability and activity allows more accurate recovery of site variability using an elastic network model (71), an idea that could be further explored using recent protein models such as LigandMPNN (72).

Our results, together with recent work on learned evolutionary trajectories (39), suggest that such simulated trajectories can also provide information for downstream evolutionary inference and prediction. They have been used to interpret directed evolution experiments (35), forecast viral protein evolution (12), and investigate site-specific evolutionary rates (33). MSA-free site-rate prediction for orphan proteins has also been explored using dedicated structure-based predictors such as EvoRator (73). Another application lies in residue–residue coupling signals extracted from simulated alignments for structural modeling, supported by experimental evolution studies showing that such co-variation signals become increasingly pronounced over successive generations and reflect structural constraints (34, 74), together with recent work showing that simulated evolutionary trajectories can improve structure prediction for evolved proteins compared with single-sequence input alone (75).

Important limitations remain. First, the fitness-based simulations inherit the limitations of the models used to define fitness: protein language models, inverse folding models, physics- and knowledge-based energy functions, DCA, and other approaches each capture only part of the biological fitness landscape. For example, structure-based inverse folding models, such as ProteinMPNN and ESM-IF, as used here, evaluate sequence compatibility against specified backbones and therefore do not explicitly account for backbone flexibility and conformational fluctuations. Moreover, the current simulations primarily evaluate the fitness effects of amino acid substitutions, individually or through simultaneous profile optimization with AF-back as implemented here; however, support for insertions and deletions remains limited, with recent studies beginning to address this gap (39, 76).

Second, the discrete simulations carried out here rely on mutation–selection trajectories in which protein models are repeatedly queried to evaluate proposed substitutions. While this approach is natural and convenient for incorporating diverse fitness proxies, it does not explicitly model the timing between substitution events. Simulations along a given phylogenetic tree require an explicit mapping of branch lengths onto the number of simulation steps, e.g., through a scaling coefficient. The continuous-time sequence evolution model recently proposed by Pagnani and Barrat-Charlaix addresses this gap for DCA-based fitness landscapes by using the Gillespie algorithm (37). Extending this formulation to other protein models could generalize continuous-time fitness-guided simulation beyond DCA.

Finally, fitness-constrained simulation introduces additional computational cost because protein models must be repeatedly evaluated along evolutionary trajectories. This cost varies across models (SI Table 1), making improved computational efficiency an important consideration for scaling the framework.

In summary, programmable fitness objectives provide a practical route for connecting molecular mechanisms to evolutionary dynamics. By making the assumed selective constraints explicit through protein models, protein evolution simulation can move beyond generating realistic sequence variation toward testing what structural or functional features are sufficient to shape the evolutionary signatures observed in natural proteins. This view complements classical evolutionary models by allowing molecular hypotheses to be encoded directly into simulated evolutionary trajectories in a flexible way.

## Materials and methods

### Dataset

We compiled a protein dataset from the PDB database with a maximum sequence identity of 30%. To ensure structural quality and computational efficiency, we filtered entries by a resolution cutoff of 2.5 Å and a sequence length between 50 and 200 amino acids. A total of 100 protein chains were randomly selected to form the dataset used throughout this study. For each protein, we retrieved multiple alignments, corresponding phylogenetic trees, and residue-level evolutionary rates from the ConSurf-DB database (45).

### Protein models used to guide the sequence simulation in this study

We employed a diverse set of structure-based, physics-based, and sequence-based protein models to evaluate sequence fitness during evolutionary simulations (see Supplementary methods for details). Specifically, we used ProteinMPNN to estimate conditional sequence likelihoods given backbone structures, ESM-IF to assess sequence– structure compatibility via inverse folding probabilities, and PyRosetta to compute mutation-induced stability changes based on an all-atom energy function. In addition, a direct coupling analysis (DCA) model trained on the multiple sequence alignment of each protein in the compiled dataset was used to assign statistical energies reflecting evolutionary constraints, and the protein language model ProGen2 was used to evaluate sequence likelihoods under learned natural protein distributions. In all cases, model-specific scores were used as proxies for sequence fitness to guide the simulations.

### Sequence simulation under the fitness constraint

To simulate forward evolution from a given protein, we employed a mutation–selection framework. At each step, a mutation was introduced by randomly selecting a position in the sequence and substituting it with a randomly chosen amino acid. The resulting mutant was then evaluated using the specified scoring model to quantify its “goodness” or energy. To convert this property into a fitness value, we applied a commonly used soft-threshold model in protein energetics:

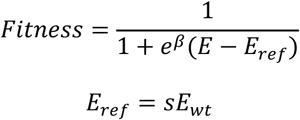

where *β* denotes the inverse temperature (1.0 used in this study), and *E* and *E_ref_* represent the energy of the current protein and the reference energy, respectively. The reference energy *E_ref_* is modulated by a scaling factor *s*, thereby controlling the stringency of selection. An *s* value of 0.5 sets the reference energy to half the wild-type energy. Thermodynamically, this fitness mapping function assumes that a protein’s fitness corresponds to the fraction of molecules occupying the ground (native) state under the equilibrium. Once the fitness of the proposed mutant was obtained, the selection was performed by comparing the new fitness *f*_j_ and the old fitness *f_i_*. Throughout this study, we adopted a birth-death model to govern the acceptance of mutations:

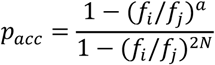

where *a* = 1 or 2 for a diploid or haploid population, respectively, and *N* represents the effective population size. The proposed mutation was accepted or rejected based on the fixation probability *p_acc_*. Following prior studies on sequence simulation (77), we set *a* = 2 and *N* = 100 in all simulations. To improve the fixation efficiency, an accelerated birth-death process was employed as previously described (77),

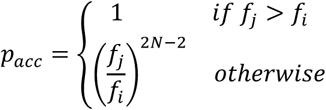

This accelerated algorithm accepts all mutations that increase fitness, thereby substantially speeding up equilibration while preserving the stationary distribution of the original process, as demonstrated in previous studies (77). The process was iterated until the specified number of substitutions had accumulated. For each protein in the compiled dataset, we performed 50 trajectories of 500 substitutions accumulated for each protein model.

### Sequence simulation under empirical substitution models

As a baseline representing traditional sequence simulation approaches, we employed the pyvolve package (version 1.1.0, https://github.com/sjspielman/pyvolve) due to its flexibility in generating customized simulations (22). The simulation process utilized a continuous-time Markov process, where substitutions between amino acids were determined by empirical substitution models (LG used throughout this study) (19). Similar to the fitness-constrained simulation, we performed 50 trajectories, each consisting of 500 substitutions, for every protein in the compiled dataset.

### AlphaFold2 backpropagation

The ColabDesign package was employed to conduct the backpropagation through the AlphaFold2 network (66) (version 1.1.3, https://github.com/sokrypton/ColabDesign). The backpropagation process was guided by a customized loss function that generates gradients to update the sequence PSSM profile. In this work, the loss function is composed of three metrics, pLDDT showing the prediction confidence, RMSD to measure the similarity between the predicted structure of the generated sequence and the starting (wildtype) structure, and a contact loss to quantify the average number of contacts one residue has with others. A weight of 0.5, 0.5, and 1.0 was assigned to the pLDDT item, the RMSD item, and the contact item, respectively. For each protein, the backpropagation was initialized with the wildtype sequence and went through 500 gradient descent steps. No structural templates were provided to AlphaFold2; the wild-type structure was used only as the optimization target. A learning rate of 0.1 was employed with the Stochastic gradient descent (SGD) optimizer. To prevent the simulation from being a simple optimization procedure, we implemented a dynamic threshold-based loss function to mimic neutral evolution,

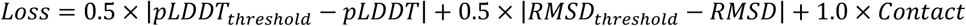

Here, *pLDDT_t_*_ℎ*res*ℎ*old*_ is a target confidence score ranging from 0.9 to 1.0, and *RMSD_t_*_ℎ*res*ℎ*old*_ is a structural deviation threshold ranging from 0.0 to 0.2. A *pLDDT_t_*_ℎ*res*ℎ*old*_ and an *RMSD_t_*_ℎ*res*ℎ*old*_ were randomly sampled for each backpropagation step.

### Sequence simulation based on given phylogeny

For the discrete simulation schemes, we performed forward simulations along the phylogenetic tree for each protein, using the wild-type sequence as the root sequence. For the AlphaFold2 backpropagation, the simulation along the tree was conducted in a similar way: the protein structure and the corresponding wildtype sequence was used as the root sequence. For each branch connecting two nodes (internal nodes or leaves), a single AlphaFold2 backpropagation was performed in a number of specific steps. For each branch of the discrete simulation, the number of simulation steps was determined by multiplying the phylogenetic branch length and the protein length. For the AF2 backpropagation, the branch length was instead multiplied by a fixed scaling factor of 200 steps per phylogenetic unit. A minimum of one simulation step was used for each branch, such that longer branches underwent proportionally more simulation steps.

### Comparison of amino acid distribution

Following previous studies (23), we quantified the similarity between the simulated trajectories and the natural multiple alignments by computing the sum of the Kullback-Leibler (KL) divergence across all protein site:

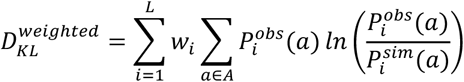

where *a* denotes a specific amino acid type from the set of 20 standard residues *A* and *i* refers to a site in a protein sequence of length *L*. The KL divergence was computed at each site *i* and weighted by the fraction of aligned (non-gap) residues in column *i* of the multiple sequence alignment. The weighted divergences were then summed across all sites to yield an overall divergence score, serving as a measure of how much the simulated trajectories deviated from natural alignments.

### Calculation of site evolutionary rates

For a given sequence evolutionary trajectory generated under the stepwise mutation framework, the site-specific evolutionary rates were estimated by recording the number of substitution events at each site throughout the simulation and using these counts as a proxy for relative evolutionary rates. For a given evolutionary trajectory generated via AlphaFold backpropagation, site-specific evolutionary rates were estimated using the aforementioned methods once the intermediate sequences, which were represented as logits over 20 amino acid types at each site, were decoded into discrete amino acid sequences by selecting the highest-logit amino acid at each position (argmax). Substitution events were then counted as changes in the argmax-decoded amino acid state between consecutive intermediate sequences.

### Estimation of substitution matrices

To compare the evolutionary dynamics produced by different fitness-guided simulations, we derived empirical amino acid substitution models directly from simulated trajectories. Substitutions observed across trajectory steps were aggregated to estimate a Markov substitution process, from which equilibrium amino acid frequencies and exchangeability parameters were inferred. The resulting rate matrices were normalized to unit expected substitution rate and compared with standard empirical models such as LG (See Supplementary methods for details).

### Residue-residue co-variation detection

The residue-residue co-variation signals encoded in sequence simulation were detected and compared at the level of pairwise coupling intensity. We simulated the phylogenetically structured alignments for each protein. For each simulated alignment, the plmDCA package (version 1.0.4, https://github.com/pagnani/PlmDCA.jl) was applied to infer the pairwise coupling *J_i_*_j_ (43). The derived couplings *J_i_*_j_ from LG-, ProteinMPNN- and AF-back-based simulations were compared with those computed from the corresponding natural alignment from ConSurf-DB (45), characterized using Spearman correlation coefficient.

### Sequence evaluation

Sequences produced along the simulation were evaluated from three aspects: naturalness of protein sequences, structure quality, and energetics. ESM-2 (version 1.0.3, https://github.com/facebookresearch/esm) is a protein language model that predicts masked positions in a sequence (78). The pseudo-likelihood *L_ESM_*_2_(*S*) of a sequence *S* = (*s*_1_, *s*_2_, …, *s_L_*) is calculated by masking each amino acid *s_i_* and estimating the conditional probability of *s_i_* given the remaining sequence *S*\*s_i_* (See Supplementary methods for details). ESMFold (version 1.0.3, https://github.com/facebookresearch/esm) was employed to predict the structures of evolved proteins in default settings. The Discrete Optimized Protein Energy (DOPE) method implemented in MODELLER (version 10.6, https://salilab.org/modeller/) was used to evaluate the energetics of evolved proteins (79). For each evolved sequence, the corresponding wild-type PDB structure was used as a template to generate three independent comparative models using the standard MODELLER AutoModel protocol, including conjugate-gradient optimization followed by simulated-annealing refinement. DOPE scores were averaged across the three models, and the wild-type sequence was evaluated using the same procedure. Each metric (ESM2 likelihood, ESMFold pLDDT, and DOPE score) was then normalized by dividing the score of the evolved sequence by that of the wild-type sequence.

### Evolving proteins under composite fitness objectives

We linearly combined fitness values derived from scores from three deep learning–based models: ProGen2, ProteinMPNN, and ESM-IF. For each combination scheme, the fitness proxies derived from different models were assigned the same weight of 1.0. We applied the same simulation procedure to all seven single- and multi-model scenarios on the 100-protein dataset, and evaluated the resulting trajectories using the methods described above.

### Evolving proteins under homodimer constraints with ProteinMPNN

We selected a set of well-characterized proteins with biologically relevant homodimeric forms as model systems, including catabolite activator protein (CAP, PDB ID: 1G6N), interleukin-8 (IL-8, 1IL8), ribonuclease III (RNase III, 1JFZ), triose phosphate isomerase (TIM, 1TIM), HIV-1 protease (2BQV), alkaline phosphatase (ALP, 3TG0), nerve growth factor (NGF, 4EC7), malate dehydrogenase (MDH, 4MDH), Cro repressor (5CRO), aminotransferase (6HNB), and estrogen receptor α (ERα, 9MDV) (80–90). All structures were processed using MODELLER to reconstruct missing residues in the deposited PDB files (79). For each protein, sequence evolution was simulated using fitness scores derived from ProteinMPNN under both monomeric and dimeric contexts. To evaluate sequences in the dimer context, the full dimer structure was provided as input to ProteinMPNN, and sequence log-probability profiles were computed by combining the profiles from both chains. A range of weighting schemes was explored (0 ≤ *w*_1_ ≤ 1, with *w*_1_ + *w*_2_ = 1). For each condition, we generated 100 evolutionary trajectories, each comprising 500 accepted substitutions. The resulting trajectories were analyzed using the methods described above. Interacting residues were identified using the PDBePISA web server (https://www.ebi.ac.uk/pdbe/pisa/) (91).

To generate decoy dimer conformations, the HDOCK web server (http://hdock.phys.hust.edu.cn/) was employed using default settings (54). For each protein in the dimer-based simulation set, HDOCK generated 100 dimeric models. These models were then evaluated using the DockQ package (https://github.com/wallnerlab/DockQ) (92). Models with interface properties comparable to those of the experimentally determined dimer structure (at least 50% of the buried surface area and 50% of the contact numbers), while adopting conformations distinct from the native structure (interface RMSD > 6 Å, ligand RMSD > 10 Å, and DockQ score < 0.1), were retained as the candidate pool. Interface area was calculated as [SASA(A) + SASA(B) − SASA(AB)] / 2 using FreeSASA (93), and interfacial contacts were defined as unique residue–residue pairs with at least one heavy-atom pair within 5.0 Å between chains A and B. When these criteria yielded an insufficient number of candidates, the selection thresholds were progressively relaxed until a sufficient number of decoy models was obtained, while prioritizing models that most closely satisfied the original criteria. Three decoys were then randomly selected from the resulting candidate pool and subjected to the same simulation and analysis procedures described above.

### Evolving metamorphic proteins under multi-conformational constraints with ProteinMPNN

We selected three well-characterized fold-switching systems reported in recent reviews (55): XCL1 (the α+β fold, PDB ID: 1J8I (56), and the all-β fold, PDB ID: 2JP1 (57)), KaiB (C-terminal βααβ fold, 2QKE (59); C-terminal αββα fold, 5JYT (60)), RfaH (C-terminal α-hairpin fold, 2OUG (61); C-terminal β-roll fold, 6C6S (62)), and included SOD1 as a non-metamorphic comparison (the β-barrel fold, 1UXM (63); the nascent highly disordered state, 2NAM (64)). For each system, the two conformational states were trimmed to match lengths. Missing residues in the PDB structures were reconstructed using MODELLER. The sequence identities between the two states for XCL1, KaiB, RfaH, and SOD1 are 100%, 95%, 100%, and 99%, respectively (see SI for sequences used). For structures determined by solution NMR and deposited as structural ensembles in the PDB (1J8I, 2JP1, 5JYT, and 2NAM), the first model was used.

Sequence evolution simulations were initialized from the sequence of one representative PDB structure for each system. For each protein, both conformations were provided as inputs to the ProteinMPNN-based simulation framework. Where applicable, the homo-oligomeric state of the protein was maintained as represented in the corresponding PDB structure (2JP1 as a dimer, 2QKE as a tetramer, and 1UXM as a dimer). The fitness function was defined as a weighted linear combination of two fitness terms, each corresponding to one conformation. A range of weighting schemes was explored (0 ≤ *w*_1_ ≤ 1, with *w*_1_ + *w*_2_ = 1). For each weighting condition, 100 evolutionary trajectories were generated, each consisting of 500 accepted substitutions. The resulting trajectories were analyzed using the methods described above.

As a control, same-fold structural decoys were generated for one conformation of each of the three metamorphic proteins using the NormalModeRelax protocol in Rosetta (28). One hundred structural variants were generated from an experimentally determined conformation (1J8I, 2QKE, or 2OUG), and three decoys with Rosetta scores at least half as favorable as that of the corresponding experimental structure and a Cα RMSD of at least 1.5 Å were randomly selected for subsequent analysis. Each decoy was paired with the reference conformation to define a two-structure ProteinMPNN fitness objective, and evolutionary simulations were performed using the same weighting schemes and analysis procedures as for the experimentally characterized conformation pairs.

## Data availability

Source codes of SeqSimFit are available on the GitHub repository https://github.com/KYQiu21/SeqSimFit. The Colab Notebook for running the simulation schemes described in this manuscript is available at https://colab.research.google.com/drive/1DCDmWv99yDNn1it0vc_cwNzru5DqK_Pp.

## Acknowledgements

Computations were performed on the MPI-BIO cluster and the HPC system Raven at the Max Planck Computing and Data Facility. K.Q. would like to thank the IMPRS From Molecules to Organisms PhD program. K.Q. also thanks Natalie Ditz for assistance with visualization. This work was supported by institutional funds of the Max Planck Society. S.D.-H. was additionally supported by the National Science Centre (grant 2020/37/B/NZ2/03268).

