## Supplemental Materials for "Simulating protein evolution with programmable fitness objectives derived from modern protein models"

### Supplementary methods

#### Protein models used to guide the sequence simulation in this study

In this work, we employed different protein models to evaluate the fitness of proteins and guide the evolutionary simulations.

**ProteinMPNN**: The full-protein backbone ProteinMPNN model weight trained with 0.2Å noise was used throughout this study (v_48_020). ProteinMPNN outputs residue-level sequence logits conditioned on a given protein structure. For each input sequence and the given backbone, we computed the cross-entropy between the one-hot encoded sequence and the predicted logits from the model as follows,

$$D_{CCE}\big(X, f(X)\big) = \sum_{i=1}^{L} \sum_{a \in A} -X \ln\big(f(X)_{ia}\big)$$

where $X$ and $f(X)$ denote the one-hot sequence and the predicted logits, respectively. A recent study has demonstrated that the ability of ProteinMPNN to predict folding stability improves substantially when conditioned on the ground-truth (wild-type) sequence. Therefore, we adopted the conditional cross-entropy as a proxy for sequence fitness.

**ESM-IF**: Similar to ProteinMPNN, ESM-IF is an inverse folding model that takes a protein backbone structure as input and outputs sequence logits constrained by the structural context. These logits can be used to assess how well a given sequence conforms to the backbone. Following a recent study, we evaluated the likelihood of each residue in the input sequence by applying a $Softmax$ transformation to the logits and extracting the corresponding probabilities. The overall sequence likelihood was then computed by summing the likelihood scores across all residues, and this value was used as a proxy to guide the simulation.

**Rosetta**: As a physics-based approach, PyRosetta was employed to assess the stability of protein mutants. For each protein subject to simulation, the input structure was first refined using the $FastRelax$ protocol with the L-BFGS minimizer. To evaluate a specific mutant, the target mutation was introduced into the refined structure, followed by side-chain repacking within a 10Å radius around the mutated site. The mutated structure was subsequently scored using the beta_nov16_cart energy function as ΔG to represent the resulting fitness.

**Direct coupling analysis model**: For each protein in the compiled dataset, we used the plmDCA package to fit a direct coupling analysis (DCA) model based on pseudo-likelihood maximization, with regularization parameters $\lambda_J = 0.01$ and $\lambda_H = 0.1$. We utilized MSAs from the ConSurf-DB database for model training. Once trained, an input sequence with the same length as the trained alignment can be assigned with a statistical energy score representing the goodness of this sequence for the fit model.

**ProGen2**: ProGen2 is a transformer-based protein language model trained on large-scale protein sequence datasets using an autoregressive objective. The sequence likelihoods derived from ProGen2 reflect how well a given sequence aligns with the learned semantic patterns of natural proteins. We computed the log-likelihood of a protein sequence $x = (x_1, x_2, \dots, x_L)$, where each $x_i$ denotes an amino acid token, under the ProGen2 model using the standard autoregressive formulation,

$$L_{ProGen2}(S) = \sum_{i=1}^{L} \log p(x_i \mid x_1, x_2, \dots, x_{i-1})$$

at each position $i$, the model predicts a probability distribution over the vocabulary conditioned on the preceding tokens. The log-probability of the ground-truth token $x_i$ was then extracted and summed over all positions. To obtain a more robust estimate of sequence fitness, we reported the mean negative log-likelihood averaged over both the left-to-right and right-to-left directions.

**Estimation of substitution matrices**

To infer empirical amino acid substitution models from simulated evolutionary trajectories, we estimated a continuous-time Markov substitution process directly from observed mutation events across trajectories.

For each amino acid pair $a_i \to a_j$, we counted the total number of observed substitutions $N_{ij}$ between successive sequence states throughout all trajectories. In addition, for each amino acid $a_i$, we recorded the total state occupancy time $S_i$, corresponding to the cumulative number of sequence positions occupied by $a_i$ across all trajectory steps.

The instantaneous substitution rate matrix $Q$ was then estimated using the maximum-likelihood estimator for continuous-time Markov chains:

$$q_{ij} = \frac{N_{ij}}{S_i}, i \neq j$$

Diagonal elements were defined as:

$$q_{ii} = -\sum_{j \neq i} q_{ij}$$

to ensure that each row sums to zero.

Equilibrium amino acid frequencies were estimated from the normalized state occupancies:

$$\pi_i = \frac{S_i}{\sum_k S_k}$$

Finally, the rate matrix was normalized such that the expected substitution rate under equilibrium equals 1:

$$Q \leftarrow \frac{Q}{-\sum_i \pi_i q_{ii}}$$

For comparison with standard reversible amino acid models (e.g., LG), we additionally derived a symmetric exchangeability matrix $R$ using:

$$r_{ij} = \frac{1}{2}(\frac{q_{ij}}{\pi_j} + \frac{q_{ji}}{\pi_i})$$

which enforces symmetry of exchangeabilities while preserving the estimated equilibrium frequencies.

**Sequence evaluation**

Sequences produced along the simulation were evaluated from three aspects: naturalness of protein sequences, structure quality, and energetics.

ESM-2 is a protein language model that predicts masked positions in a sequence (1). The pseudo-likelihood $L_{ESM2}(S)$ of a sequence $S = (s_1, s_2, \ldots, s_L)$ is calculated by masking each amino acid $s_i$ and estimating the conditional probability of $s_i$ given the remaining sequence $S \backslash s_i$:

$$P_{ESM2}(S) = \prod_{i=1}^{L} P_{ESM2}(s_i | S \backslash s_i)$$

where

$$S \backslash s_i \coloneqq (s_1, \ldots, s_{i-1}, [MASK], s_{i+1}, s_L)$$

The pseudo-likelihood is then computed as:

$$L_{ESM2}(S) = \sum_{i=1}^{L} \log P_{ESM2}(s_i | S \backslash s_i)$$

**Visualization**

Structures were visualized with PyMOL 3.0 (https://www.pymol.org/). Other figures were plotted using the Matplotlib v3.10 and seaborn v0.13 packages.

### Supplementary figures

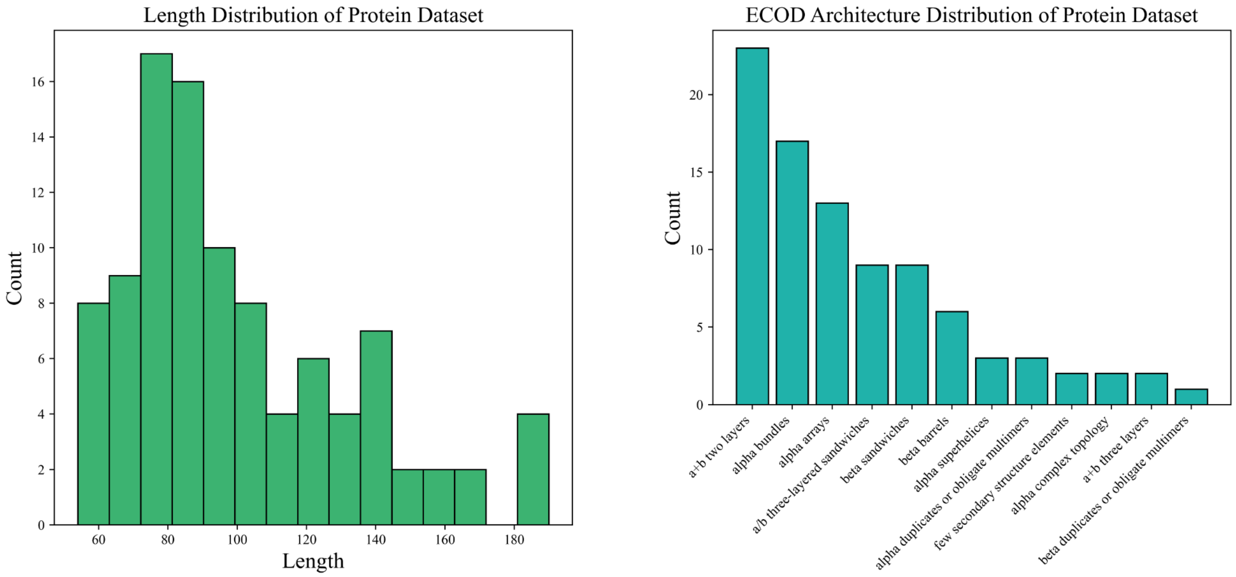


**Supplementary Figure 1: Composition of the 100-protein benchmark dataset.** (Left) Distribution of sequence lengths for the proteins used in the general benchmark. (Right) Distribution of ECOD architecture annotations, showing that the dataset covers a diverse range of protein structural architectures while maintaining sequence lengths suitable for computationally intensive fitness-guided simulations.

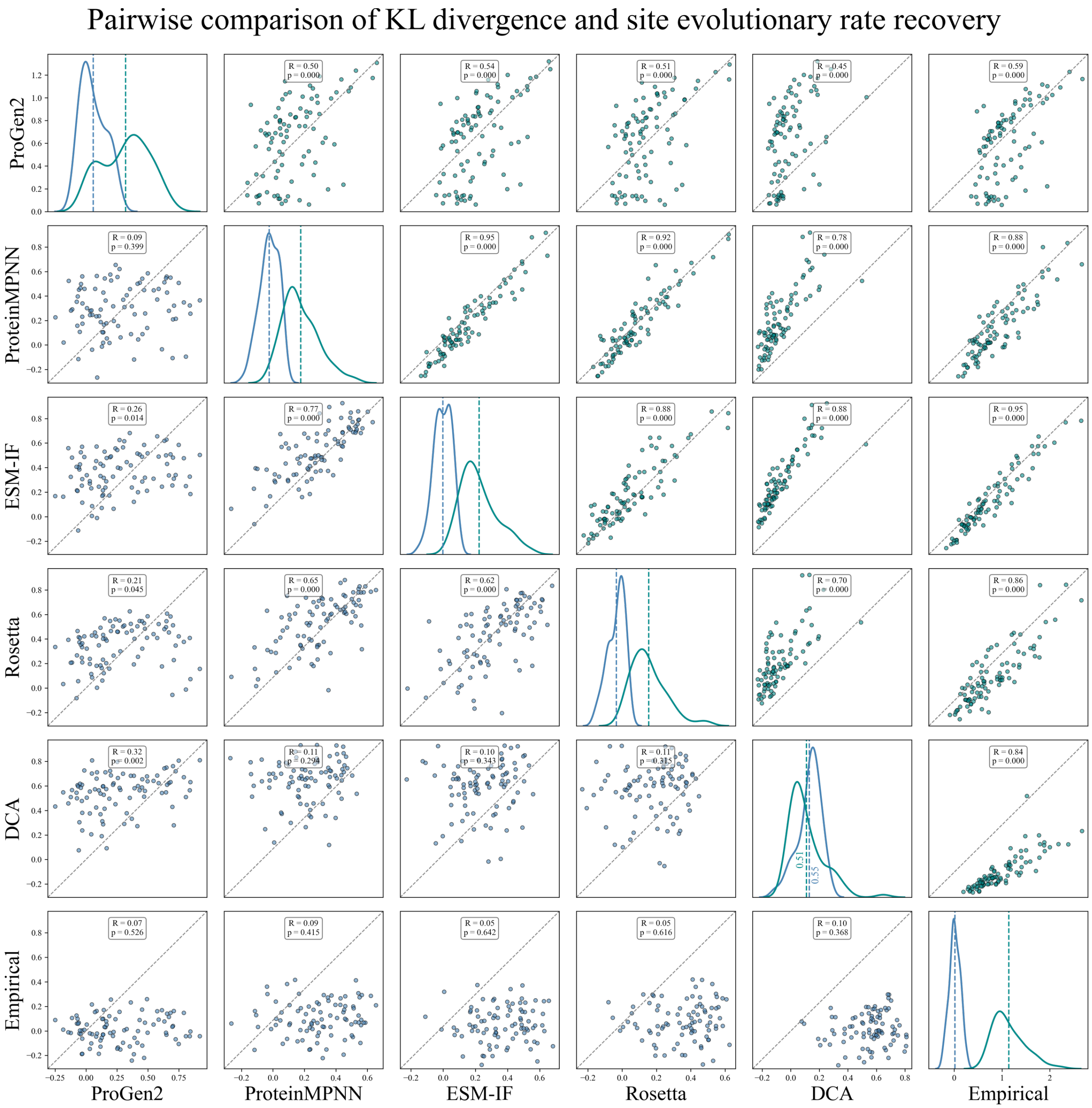


**Supplementary Figure 2: Pairwise comparison of trajectory-level metrics across simulation strategies.** Pairwise relationships between simulation methods were evaluated across the 100-protein benchmark for divergence from natural alignments and site-specific rate recovery. The upper triangle (green points) shows pairwise comparisons of per-protein site-wise KL divergence scores, each computed between simulated and corresponding natural amino acid distributions, whereas the lower triangle (blue points) shows pairwise comparisons of per-protein correlations between simulated site-specific rates and ConSurf-derived rates. Each point represents one protein, and the dashed diagonal indicates equal values between the two compared methods. Diagonal panels show the corresponding metric distributions for each method, with dashed vertical lines indicating mean values; green and blue represent sequence divergence and site-rate recovery, respectively. Correlation coefficients and p-values are shown for each pairwise comparison.

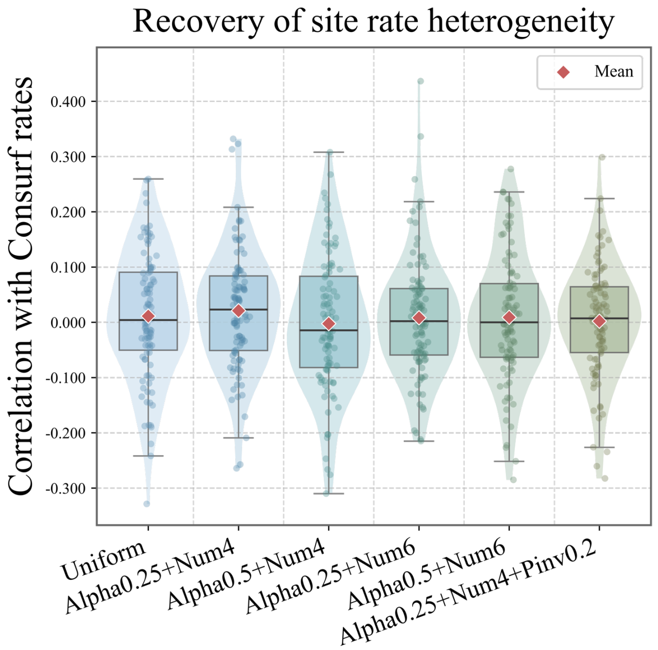


**Supplementary Figure 3: Empirical substitution models do not recover natural site-specific rate heterogeneity without alignment-derived priors.** Pyvolve simulations were performed using alternative empirical substitution and rate-model settings, including models with site-rate heterogeneity. Alpha represents the shape parameter of the discrete gamma distribution from which rates are drawn, Num represents the number of rate categories to draw, and Pinv stands for a proportion of invariant sites. For each simulation setting, site-specific evolutionary rates were estimated from accepted substitutions along simulated trajectories and compared with ConSurf-derived site rates across the 100-protein benchmark. Individual points represent proteins, red diamonds indicate means, boxes show the interquartile range, center lines indicate medians, and whiskers indicate the data range excluding outliers.

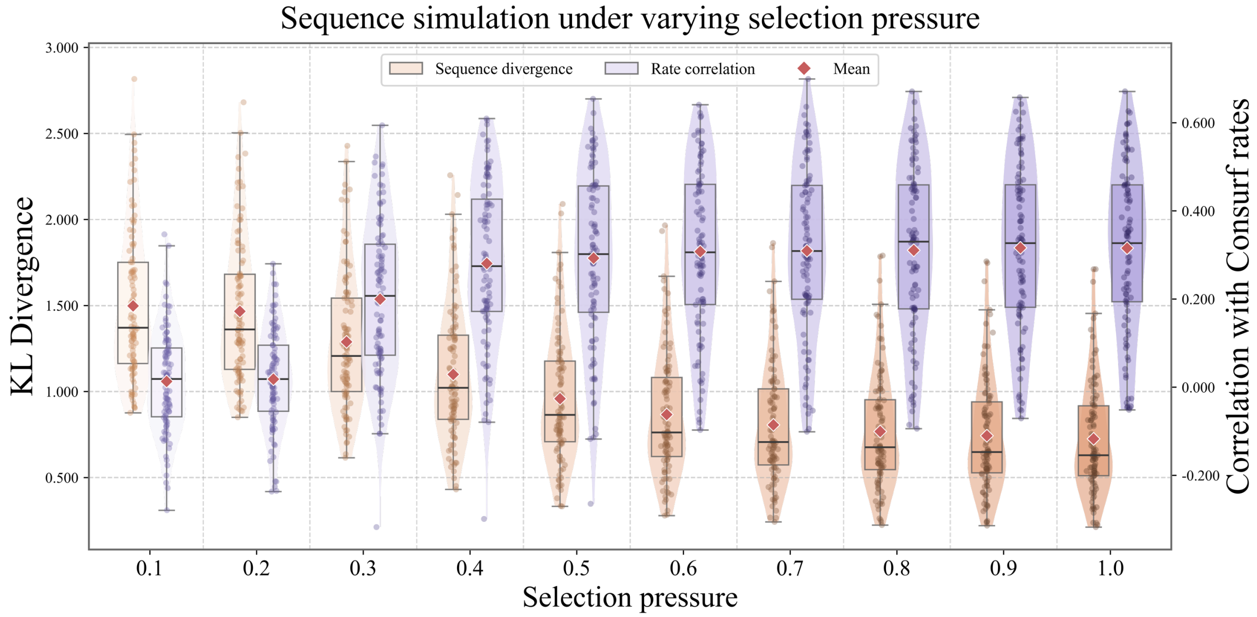


129
130 **Supplementary Figure 4: Sensitivity of ProteinMPNN-guided simulations to selection pressure.** ProteinMPNN-
131 guided simulations were repeated across the 100-protein benchmark while varying the selection pressure parameter *s*
132 from 0.1 to 1.0. For each setting, sequence divergence from natural alignments was measured by site-wise KL
133 divergence (orange, left y-axis), and recovery of site-specific evolutionary rate heterogeneity was measured by
134 correlation with ConSurf-derived rates (purple, right y-axis). Individual points represent proteins, red diamonds
135 indicate means, boxes show the interquartile range, center lines indicate medians, and whiskers indicate the data
136 range excluding outliers.

137

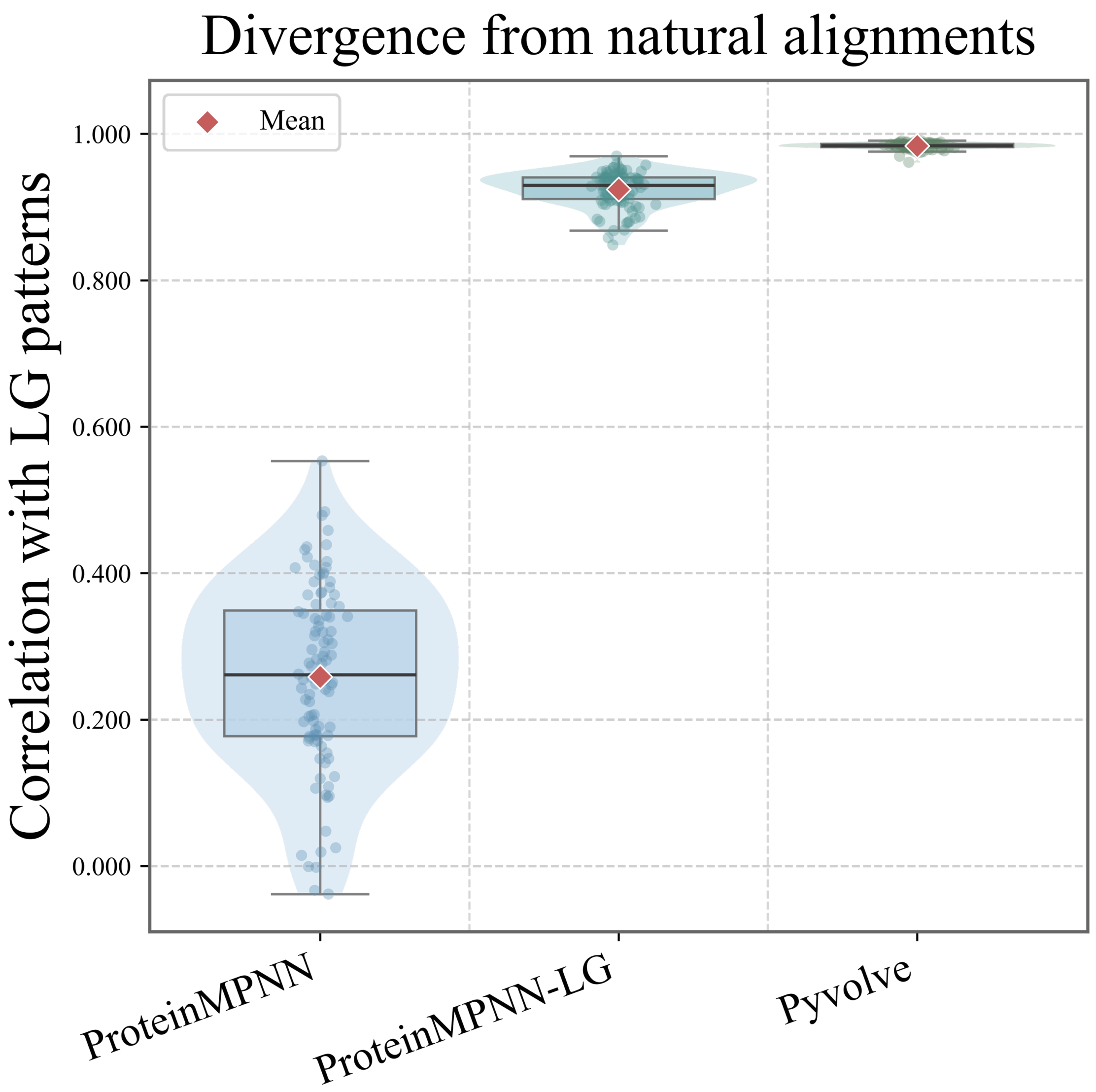


**Supplementary Figure 5: Effect of LG-based mutation proposals on substitution patterns in ProteinMPNN-guided simulations.** ProteinMPNN-guided simulations were repeated using different mutation proposal schemes, including uniform mutation proposals, LG-based mutation proposals, and Pyvolve LG simulations as an empirical substitution-model reference. Substitution matrices inferred from accepted mutations were compared with the LG exchangeability matrix. Individual points represent proteins, red diamonds indicate means, boxes show the interquartile range, center lines indicate medians, and whiskers indicate the data range excluding outliers.

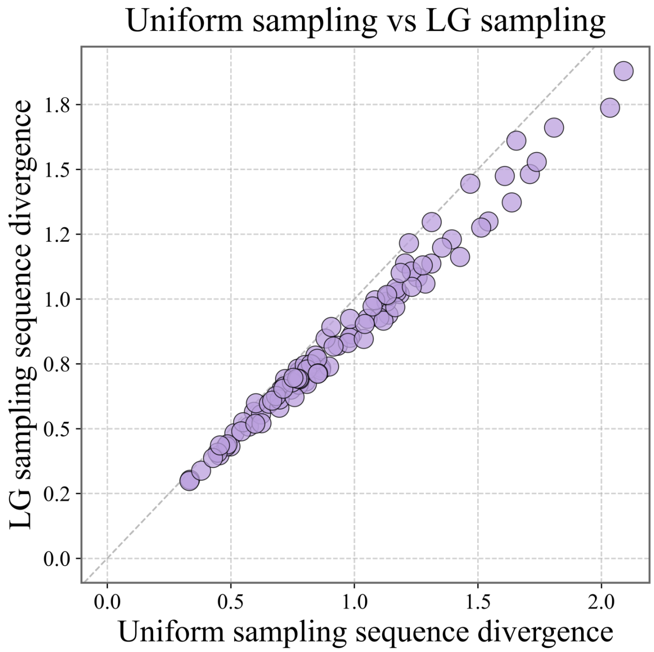


**Supplementary Figure 6: LG-based mutation proposals preserve the improved sequence divergence of ProteinMPNN-guided simulations.** ProteinMPNN-guided simulations using LG-based mutation proposals were compared with standard ProteinMPNN-guided simulations using uniform mutation proposals across the 100-protein benchmark. Each point represents one protein, and the dashed diagonal indicates equal KL divergence between the two proposal schemes. Most points lie close to or below the diagonal, indicating that changing the mutation proposal distribution to LG does not compromise, and in many cases further reduces, divergence from natural alignments under the same ProteinMPNN-based fitness constraint.

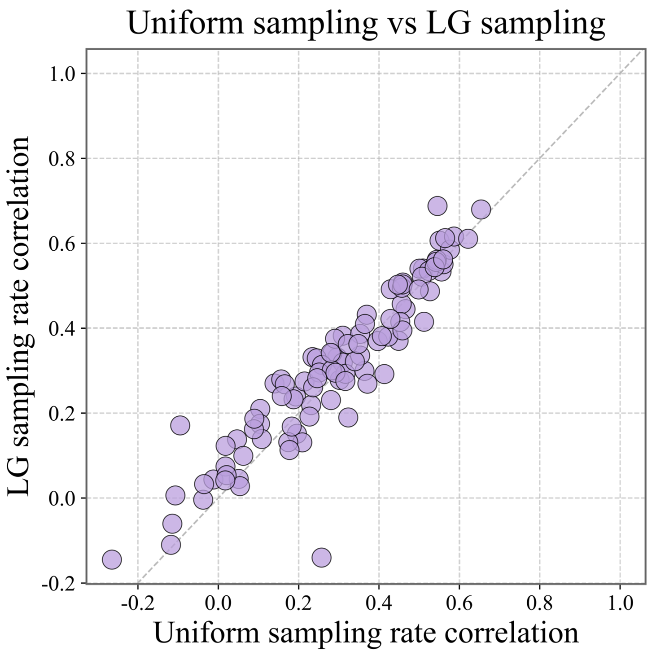


**Supplementary Figure 7: LG-based mutation proposals preserve site-rate recovery in ProteinMPNN-guided simulations.** ProteinMPNN-guided simulations using LG-based mutation proposals were compared with standard ProteinMPNN-guided simulations using uniform mutation proposals across the 100-protein benchmark. Each point represents one protein, and the dashed diagonal indicates equal recovery of site-specific evolutionary rate heterogeneity between the two proposal schemes.

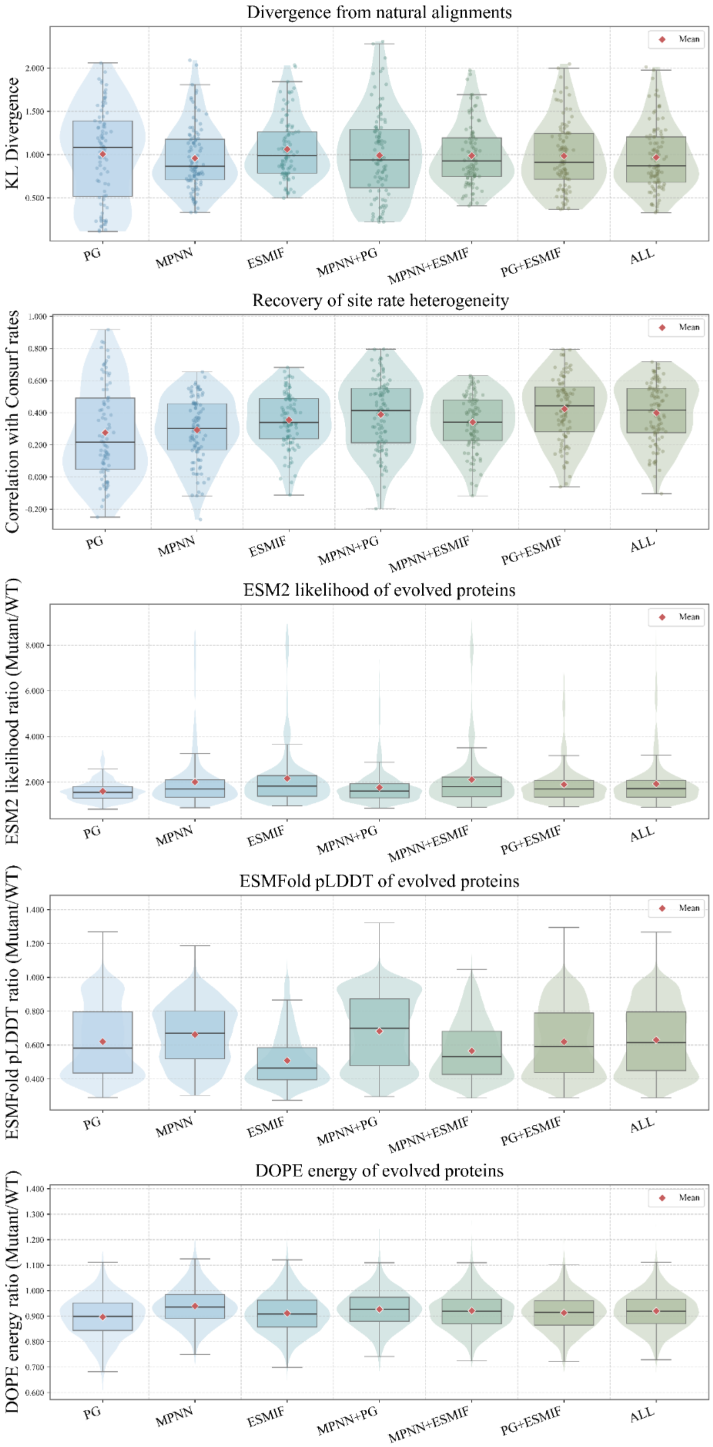


**Supplementary Figure 8: Raw distributions of evolutionary and molecular evaluation metrics for single and composite fitness objectives.** Single-model objectives based on ProGen2 (PG), ProteinMPNN (MPNN), and ESM-IF, as well as their pairwise and three-way combinations, were evaluated across the 100-protein benchmark. From top to bottom, panels show divergence from natural alignments measured by site-wise KL divergence, recovery of site-specific evolutionary rate heterogeneity measured by correlation with ConSurf-derived rates, ESM2 likelihood ratio, ESMFold pLDDT ratio, and DOPE energy ratio of evolved sequences relative to wild-type proteins. These raw distributions underlie the Z-score-normalized heatmap shown in Figure 4. Individual points represent proteins, red diamonds indicate means, boxes show the interquartile range, center lines indicate medians, and whiskers indicate the data range excluding outliers. PG: ProGen2; MPNN: ProteinMPNN; ESMIF: ESM-IF; ALL: ProGen2+ProteinMPNN+ESM-IF.

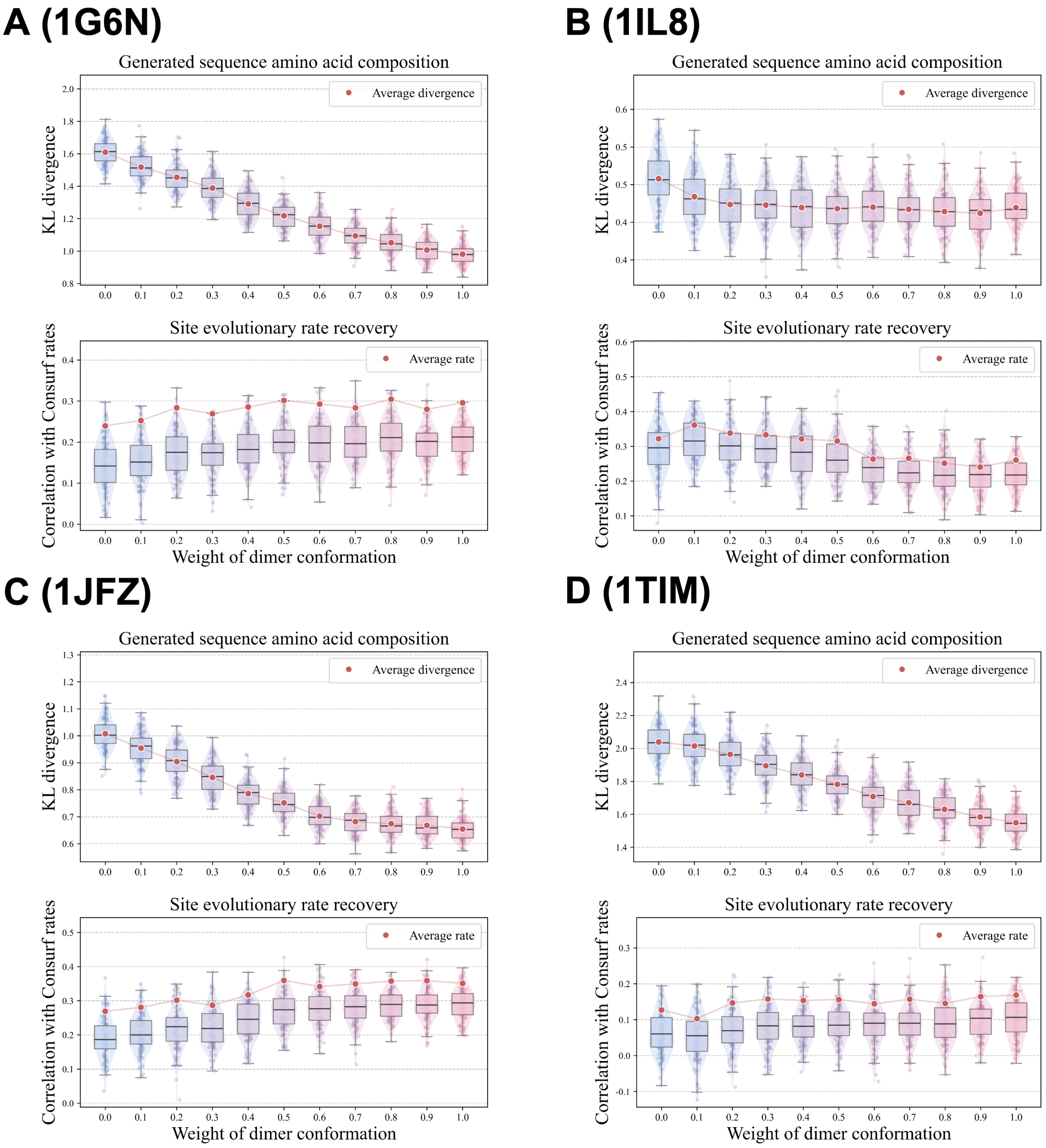

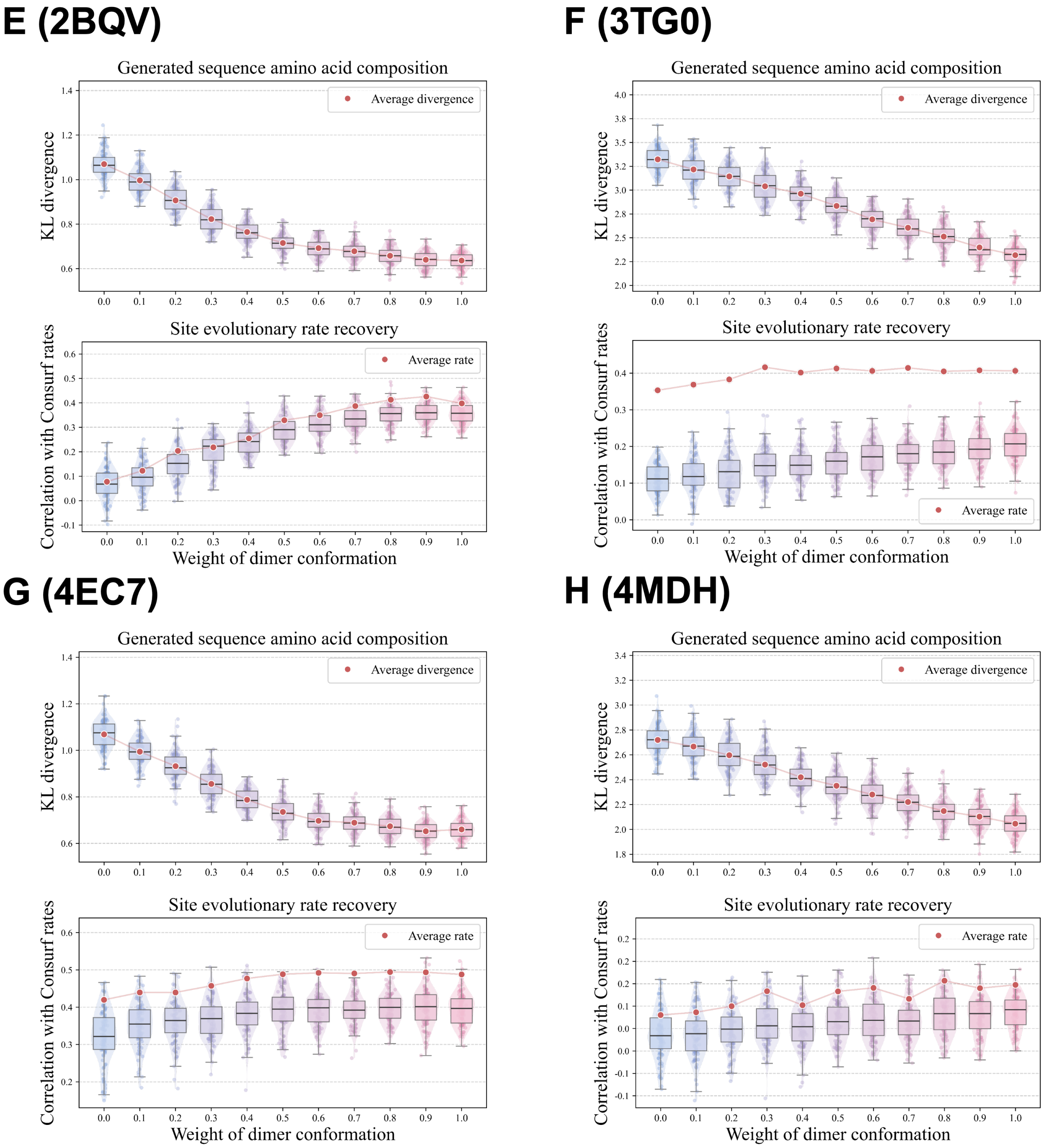
E (2BQV)
F (3TG0)
G (4EC7)
H (4MDH)
Generated sequence amino acid composition
Average divergence
KL divergence
Site evolutionary rate recovery
Average rate
Correlation with Consurf rates
Weight of dimer conformation

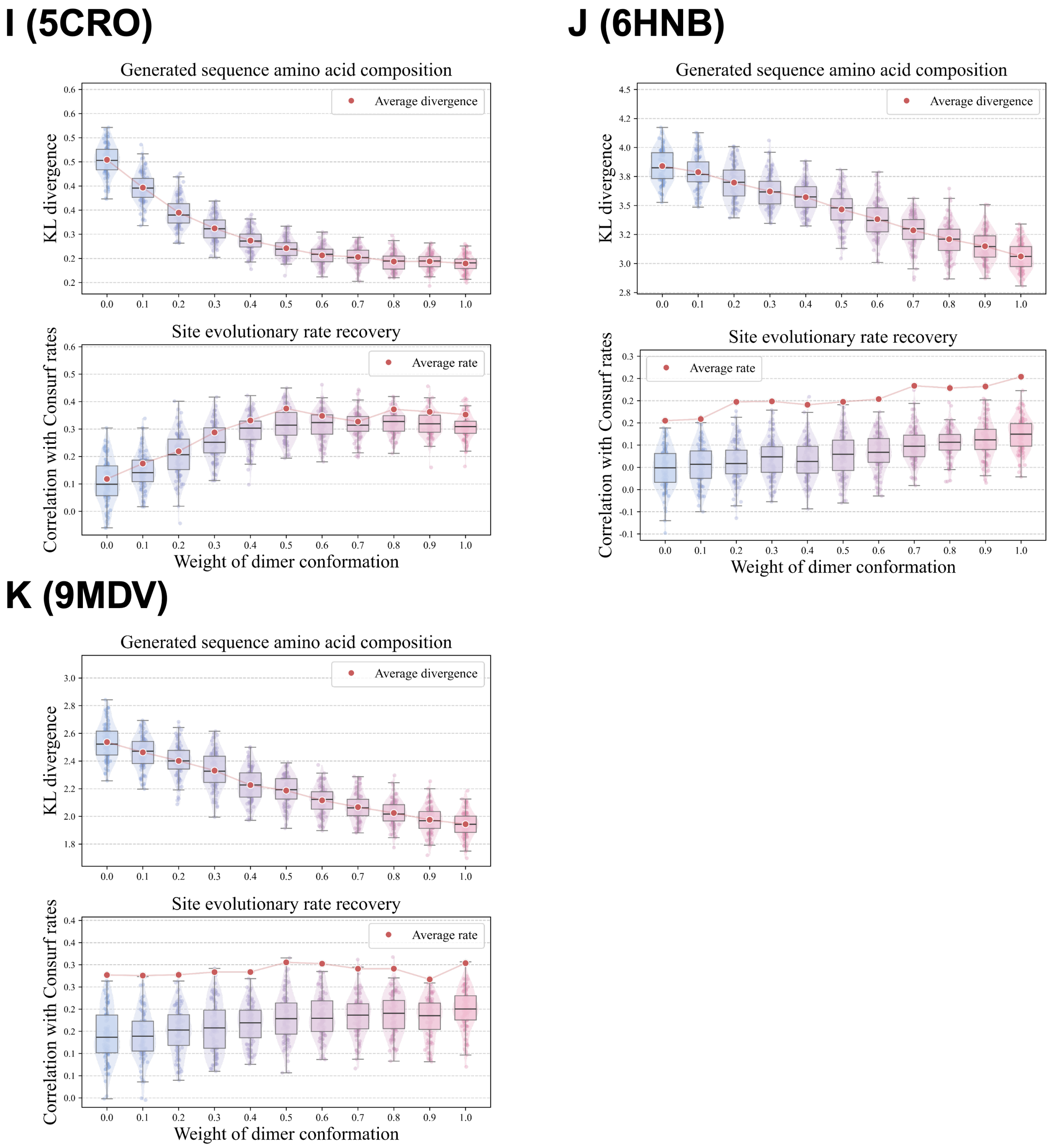


**Supplementary Figure 9: Encoding oligomeric context modulates sequence divergence and site-rate recovery in homodimeric proteins.** For each protein, the monomeric and dimeric structural contexts were used to define a weighted ProteinMPNN-based fitness objective, and simulations were performed across a range of relative weights assigned to the two conformations ($w_a + w_b = 1$). Each protein shown in Figure 5 is represented by a separate panel. In each panel, the upper plot shows the divergence between simulated trajectories and natural sequence alignments, quantified by the site-wise KL divergence between amino acid distributions. The lower plot shows the recovery of site-specific evolutionary rate heterogeneity, quantified by the correlation between simulated site substitution counts and ConSurf-derived evolutionary rates.

**A (1G6N)**

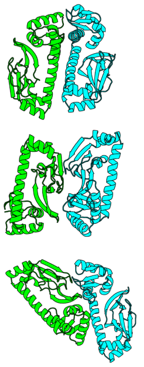

| Decoy | 1 | 2 | 3 |
|---|---|---|---|
| DockQ | 0.035 | 0.012 | 0.013 |
| iRMSD | 11.403 | 22.854 | 20.251 |
| LRMSD | 27.518 | 46.061 | 45.056 |
| BSA ratio | 0.646 | 0.692 | 0.652 |
| Contact ratio | 0.449 | 0.714 | 0.776 |

**B (1IL8)**

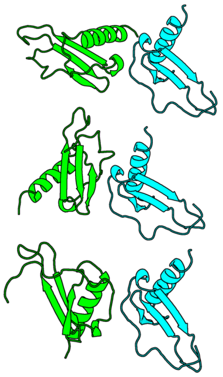

| Decoy | 1 | 2 | 3 |
|---|---|---|---|
| DockQ | 0.025 | 0.021 | 0.019 |
| iRMSD | 15.057 | 14.918 | 15.671 |
| LRMSD | 32.267 | 36.407 | 38.02 |
| BSA ratio | 0.732 | 0.717 | 0.601 |
| Contact ratio | 1.074 | 1.407 | 0.963 |

**C (1JFZ)**

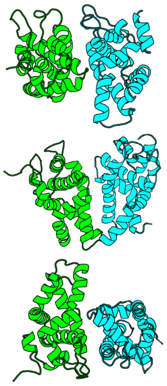

| Decoy | 1 | 2 | 3 |
|---|---|---|---|
| DockQ | 0.028 | 0.048 | 0.025 |
| iRMSD | 14.71 | 17.08 | 14.319 |
| LRMSD | 30.203 | 27.953 | 32.118 |
| BSA ratio | 0.499 | 0.675 | 0.647 |
| Contact ratio | 0.380 | 0.544 | 0.506 |

**D (1TIM)**

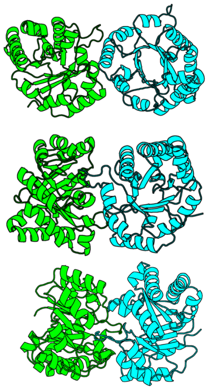

| Decoy | 1 | 2 | 3 |
|---|---|---|---|
| DockQ | 0.023 | 0.058 | 0.058 |
| iRMSD | 16.174 | 17.595 | 14.456 |
| LRMSD | 33.084 | 29.368 | 29.983 |
| BSA ratio | 0.424 | 0.498 | 0.754 |
| Contact ratio | 0.592 | 0.761 | 1.127 |

**E (2BQV)**

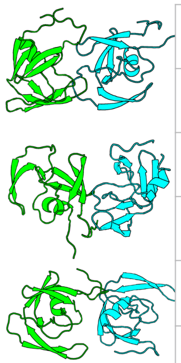

| Decoy | 1 | 2 | 3 |
|---|---|---|---|
| DockQ | 0.041 | 0.02 | 0.045 |
| iRMSD | 13.644 | 16.52 | 16.165 |
| LRMSD | 24.133 | 36.237 | 24.071 |
| BSA ratio | 0.402 | 0.407 | 0.412 |
| Contact ratio | 0.373 | 0.403 | 0.336 |

**F (3TG0)**

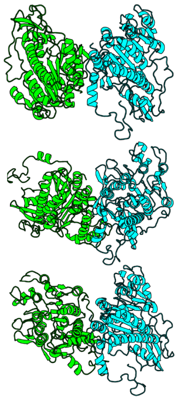

| Decoy | 1 | 2 | 3 |
|---|---|---|---|
| DockQ | 0.015 | 0.02 | 0.01 |
| iRMSD | 21.578 | 23.238 | 22.277 |
| LRMSD | 44.04 | 35.013 | 57.569 |
| BSA ratio | 0.323 | 0.355 | 0.363 |
| Contact ratio | 0.490 | 0.570 | 0.603 |

**G (4EC7)**

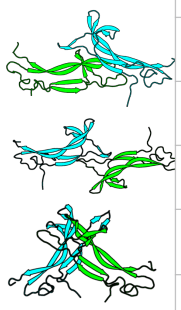

| Decoy | 1 | 2 | 3 |
|---|---|---|---|
| DockQ | 0.017 | 0.009 | 0.039 |
| iRMSD | 25.047 | 29.642 | 16.421 |
| LRMSD | 42.369 | 53.768 | 29.278 |
| BSA ratio | 0.621 | 0.489 | 0.534 |
| Contact ratio | 1.016 | 0.508 | 0.770 |

**H (4MDH)**

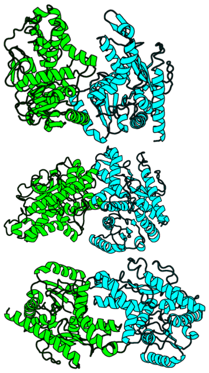

| Decoy | 1 | 2 | 3 |
|---|---|---|---|
| DockQ | 0.029 | 0.024 | 0.016 |
| iRMSD | 17.584 | 15.077 | 18.971 |
| LRMSD | 33.096 | 33.105 | 41.102 |
| BSA ratio | 0.689 | 0.628 | 0.638 |
| Contact ratio | 0.846 | 0.831 | 0.677 |

**I (5CRO)**

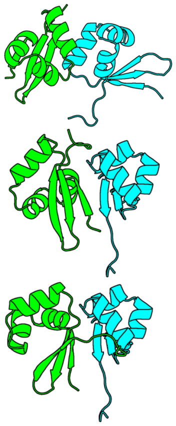

| **Decoy** | **1** | **2** | **3** |
|---|---|---|---|
| DockQ | 0.02 | 0.033 | 0.039 |
| iRMSD | 14.586 | 10.593 | 10.206 |
| LRMSD | 36.932 | 28.767 | 25.989 |
| BSA ratio | 0.800 | 0.940 | 1.072 |
| Contact ratio | 1.269 | 1.230 | 1.269 |

**J (6HNB)**

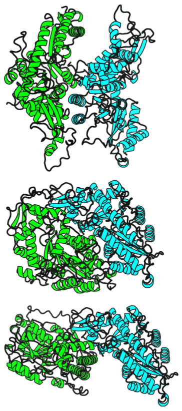

| **Decoy** | **1** | **2** | **3** |
|---|---|---|---|
| DockQ | 0.016 | 0.031 | 0.011 |
| iRMSD | 23.334 | 14.619 | 33.076 |
| LRMSD | 39.156 | 28.969 | 49.053 |
| BSA ratio | 0.317 | 0.303 | 0.316 |
| Contact ratio | 0.267 | 0.219 | 0.241 |

**K (9MDV)**

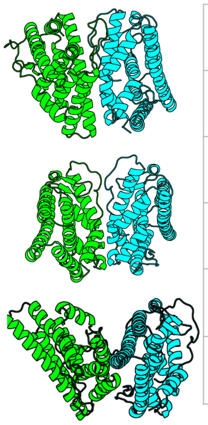

| **Decoy** | **1** | **2** | **3** |
|---|---|---|---|
| DockQ | 0.038 | 0.008 | 0.038 |
| iRMSD | 16.949 | 24.742 | 16.545 |
| LRMSD | 24.75 | 60.825 | 24.515 |
| BSA ratio | 0.715 | 0.688 | 0.738 |
| Contact ratio | 0.341 | 0.350 | 0.319 |

**Supplementary Figure 10: HDOCK generated dimer decoys.** For each protein, the conformations and statistics of the three dimer decoys were shown as an individual panel.

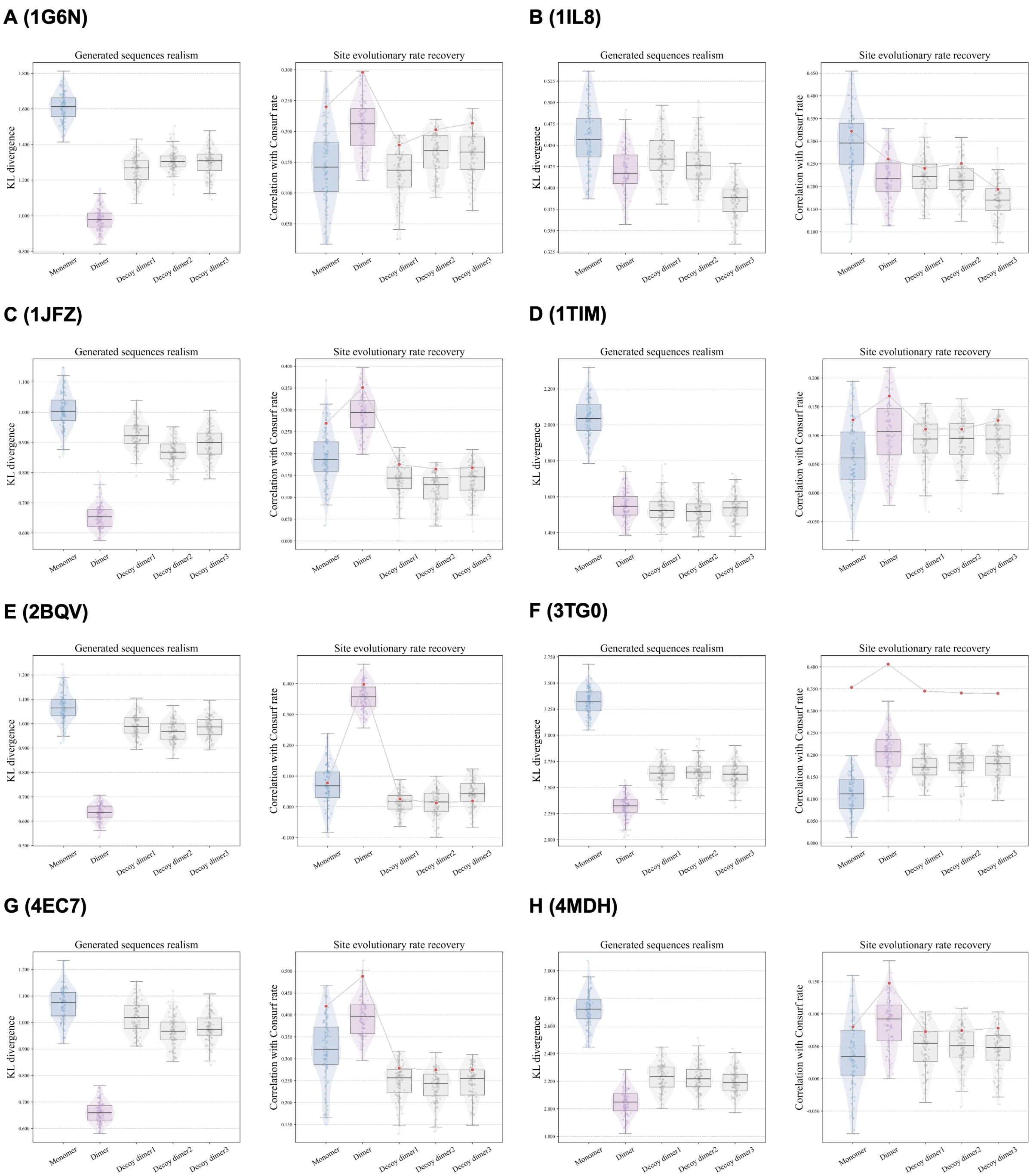

A (1G6N)
B (1IL8)
C (1JFZ)
D (1TIM)
E (2BQV)
F (3TG0)
G (4EC7)
H (4MDH)
Generated sequences realism
Site evolutionary rate recovery
KL divergence
Correlation with Consurf rate
Monomer
Dimer
Decoy dimer1
Decoy dimer2
Decoy dimer3

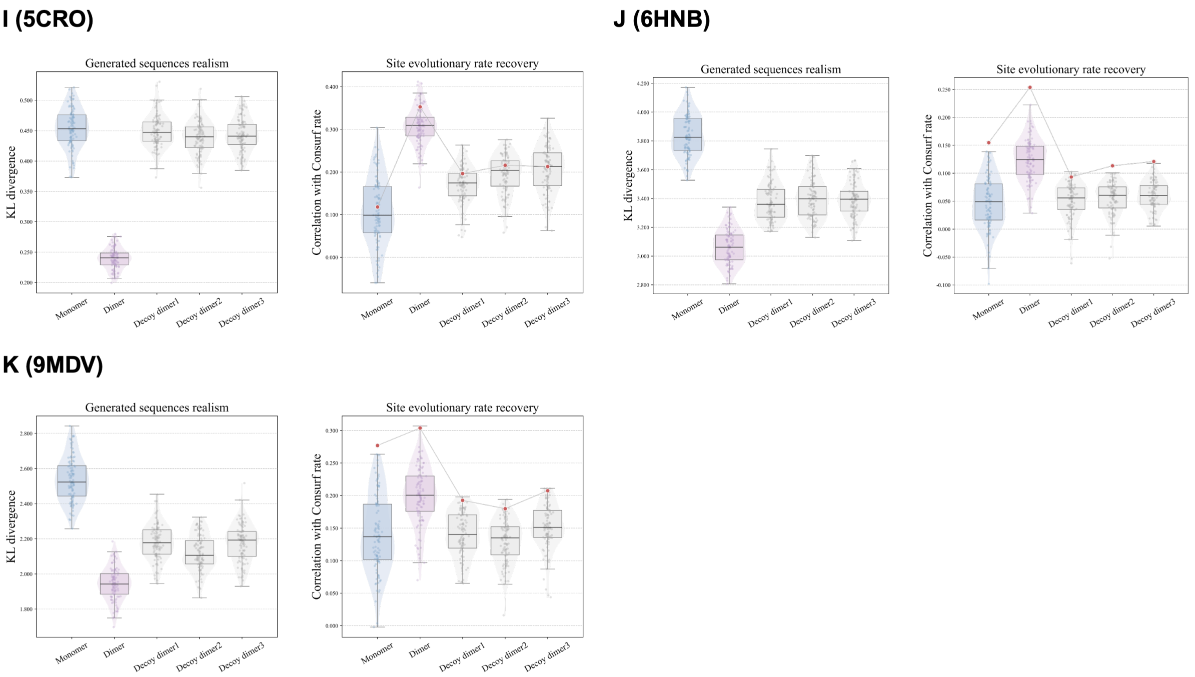


**Supplementary Figure 11: Dimeric decoys often reduce sequence divergence but fail to systematically improve site evolutionary rate recovery.** Each protein shown in Figure 5 is represented by a separate panel. In each panel, simulations constrained by the experimentally determined monomer, the experimentally determined dimer and three computationally constructed dimeric decoys were compared. The left plot shows the divergence between simulated trajectories and natural sequence alignments, quantified by the site-wise KL divergence between amino acid distributions. The right plot shows the recovery of site-specific evolutionary rate heterogeneity, quantified by the correlation between simulated site substitution counts and ConSurf-derived evolutionary rates.

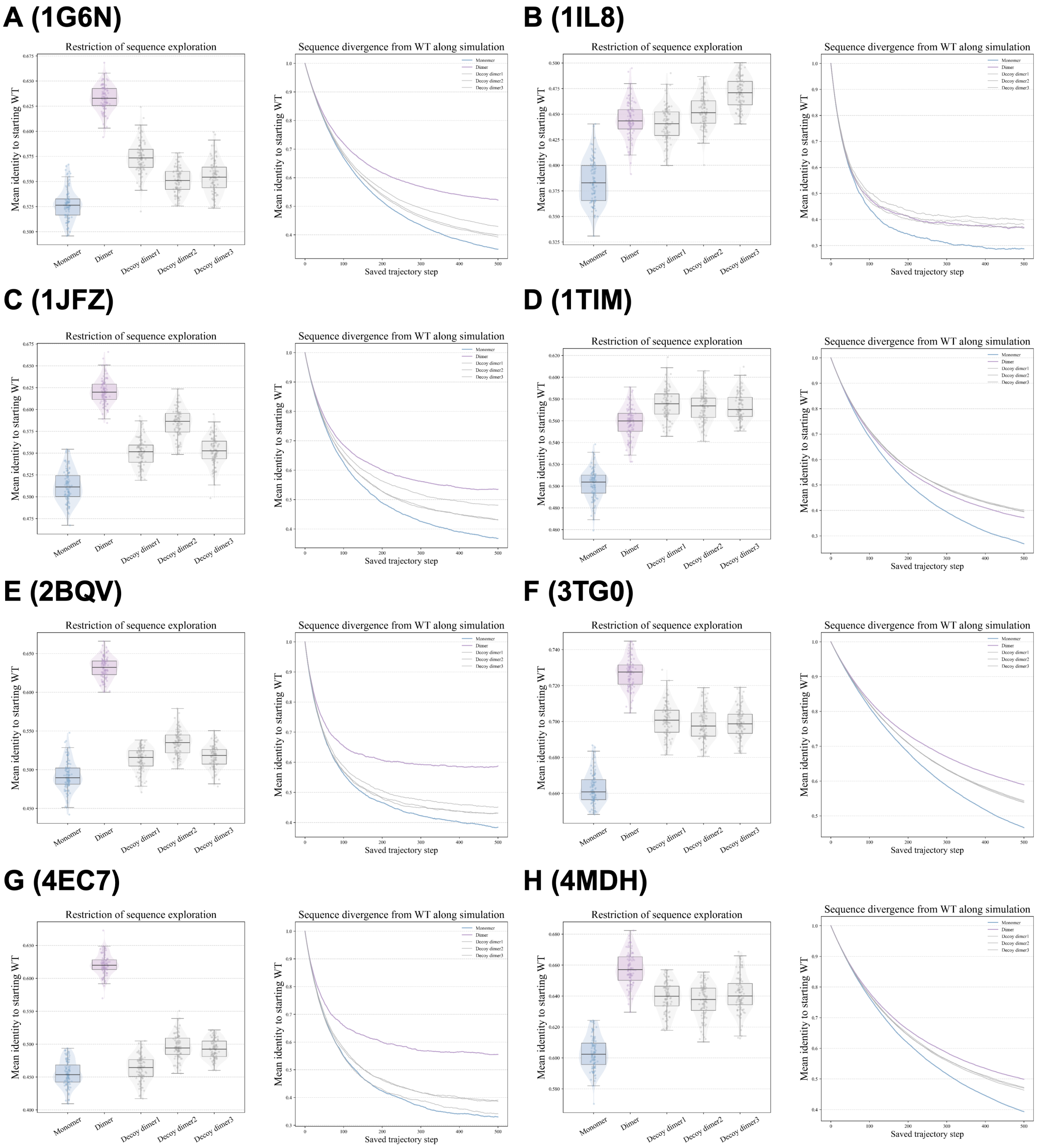
A (1G6N)
B (1IL8)
C (1JFZ)
D (1TIM)
E (2BQV)
F (3TG0)
G (4EC7)
H (4MDH)
Restriction of sequence exploration
Sequence divergence from WT along simulation
Mean identity to starting WT
Saved trajectory step
Monomer
Dimer
Decoy dimer1
Decoy dimer2
Decoy dimer3

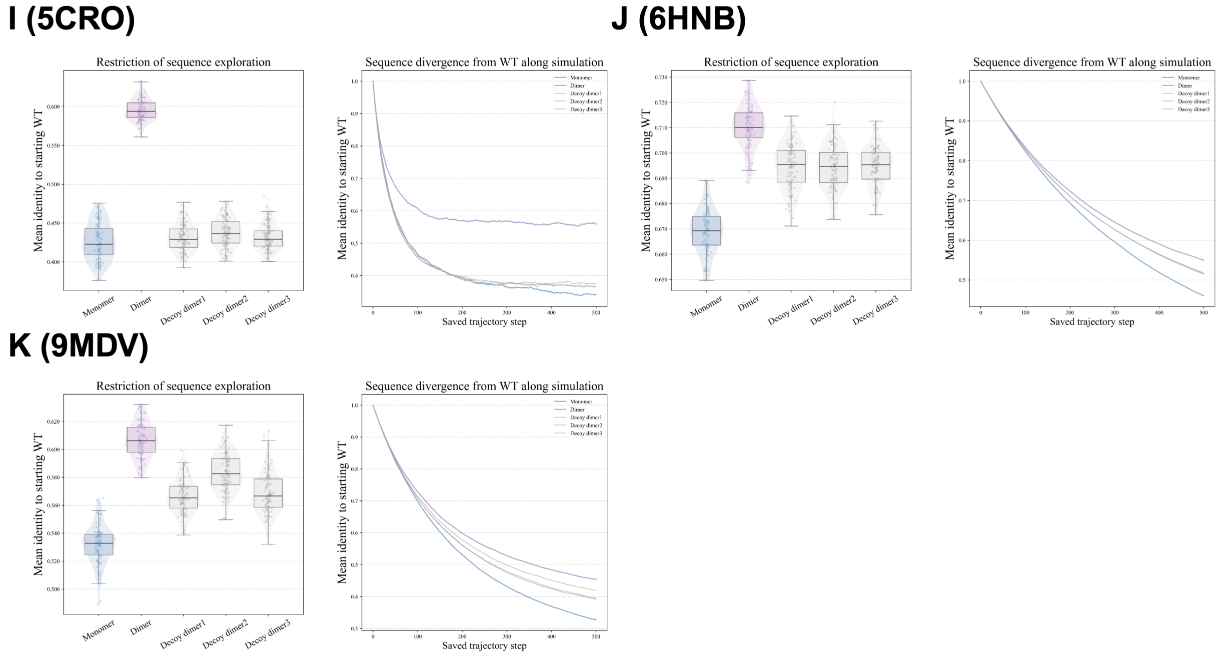


**Supplementary Figure 12: Dimeric constraints limit exploration of sequence space and reduce divergence from the wild type.** Each protein shown in Figure 5 is represented by a separate panel. In each panel, simulations constrained by the monomeric chain extracted from the experimental dimer structure, the experimental dimer and three computationally constructed dimeric decoys were compared. The left plot shows the sequence identity relative to the starting wild-type sequences. The right plot shows the change in sequence identity relative to the starting wild-type sequences over the course of the simulation.

A (1J8I)

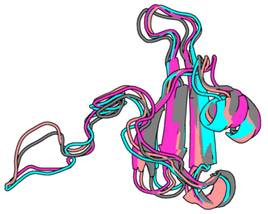

| Decoy | RMSD (Å) |
|---|---|
| 1 | 1.68 |
| 2 | 1.42 |
| 3 | 0.88 |

B (2QKE)

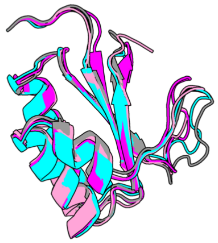

| Decoy | RMSD (Å) |
|---|---|
| 1 | 1.82 |
| 2 | 1.70 |
| 3 | 1.59 |

C (2OUG)

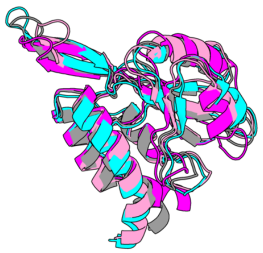

| Decoy | RMSD (Å) |
|---|---|
| 1 | 2.28 |
| 2 | 2.32 |
| 3 | 2.30 |

**Supplementary Figure 13: Rosetta generated decoys for metamorphic proteins as a second conformation.** For each protein, the conformations and statistics of the three decoys were shown as an individual panel.

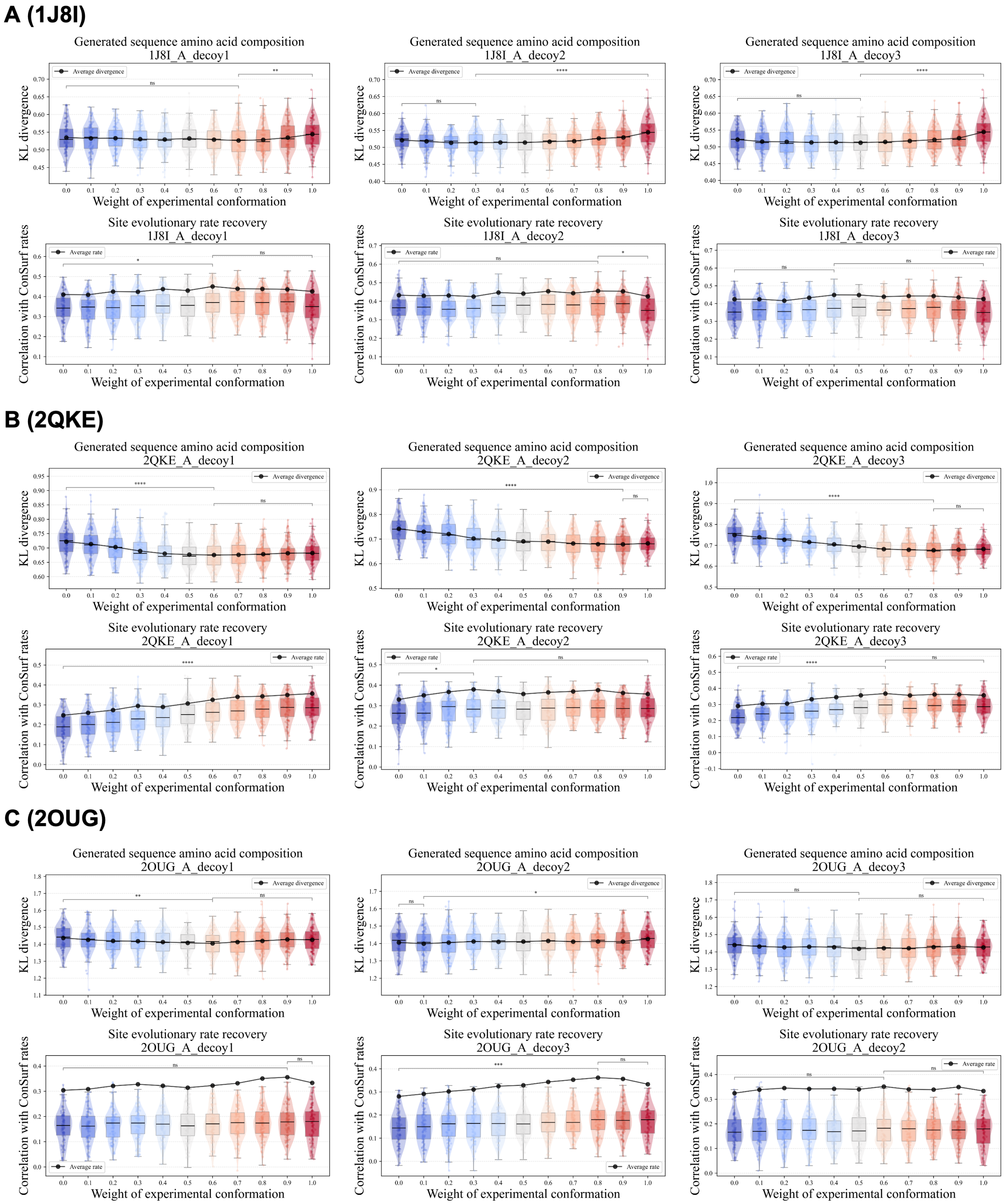


**Supplementary Figure 14: Incorporating a backbone variant into the simulation produces less pronounced valley-shaped behavior in sequence divergence and weaker peaks in site rate recovery.** For each protein, one experimentally characterized conformational state and three independently generated same-fold backbone variants obtained using Rosetta were used to define weighted ProteinMPNN-based fitness objectives, and simulations were performed across different relative weights assigned to the experimental structure and each decoy ($w_a + w_b = 1$). (A) 1J8I, (B) 2QKE, and (C) 2OUG.

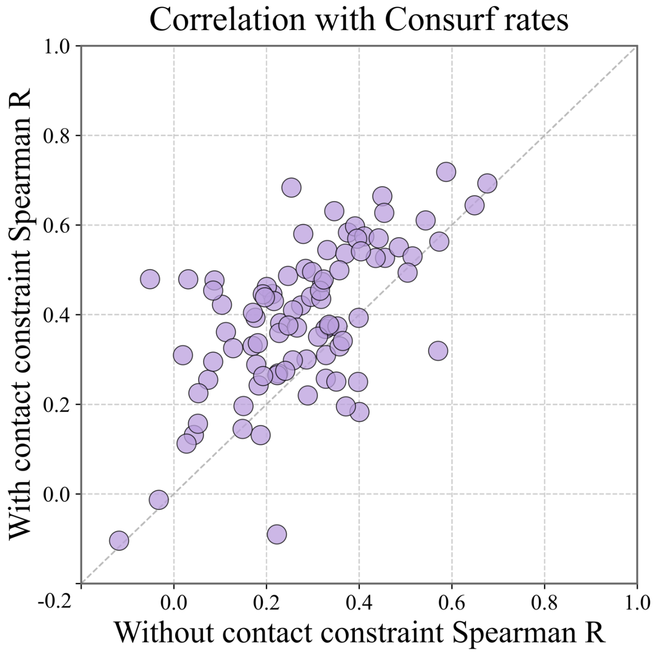


**Supplementary Figure 15: Contribution of the contact loss term to site-rate recovery in AlphaFold-backpropagation trajectories.** AF-back simulations were performed with and without the contact term in the structural loss function and compared across the 100-protein benchmark. Each point represents one protein, and the dashed diagonal indicates equal recovery of site-specific evolutionary rate heterogeneity between the two settings. Points above the diagonal indicate stronger agreement with ConSurf-derived site rates when the contact loss term is included. The predominance of points above the diagonal indicates that the contact term contributes to the improved recovery of natural site-rate heterogeneity in AF-back simulations.

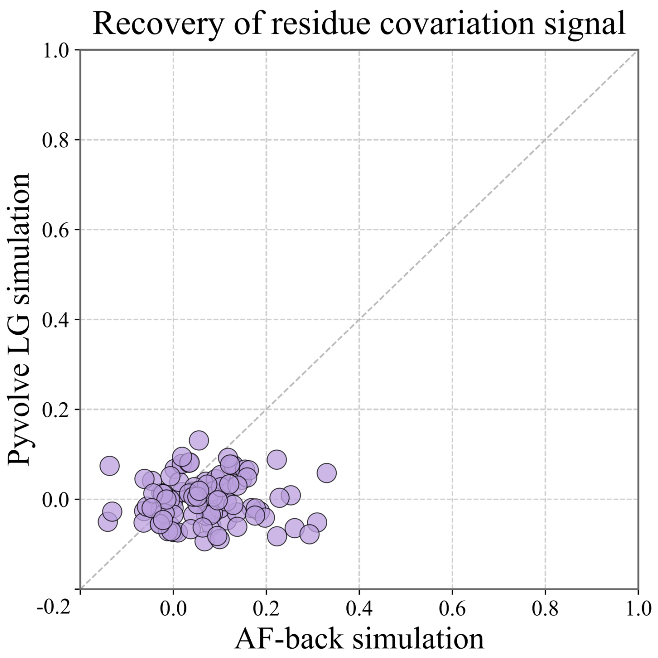


**Supplementary Figure 16 AlphaFold-backpropagation trajectories recover stronger residue–residue co-variation signals than LG simulations.** Phylogenetically structured alignments generated by AF-back were compared with alignments generated under the Pyvolve LG model across the 100-protein benchmark. Residue–residue couplings were inferred from each simulated alignment and compared with couplings inferred from the corresponding natural alignment. Each point represents one protein, and the dashed diagonal indicates equal recovery of natural co-variation signals between the two simulation strategies. Points below the diagonal indicate stronger co-variation recovery under AF-back simulation.

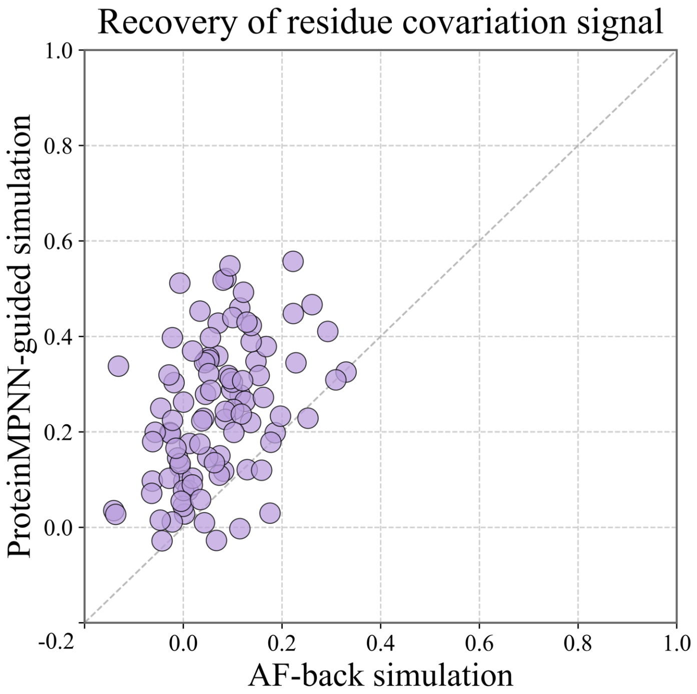


**Supplementary Figure 17: Comparison of residue–residue co-variation recovery between AlphaFold-backpropagation and ProteinMPNN-guided simulations.** Phylogenetically structured alignments generated by AF-back were compared with those generated under ProteinMPNN-guided simulation across the 100-protein benchmark. Residue–residue couplings were inferred from each simulated alignment and compared with couplings inferred from the corresponding natural alignment. Each point represents one protein, and the dashed diagonal indicates equal recovery of natural co-variation signals between the two simulation strategies. Points above the diagonal indicate stronger co-variation recovery under ProteinMPNN-guided simulation, whereas points below the diagonal indicate stronger recovery under AF-back simulation.

### Supplementary tables

| Model | Runtime (s) |
|---|---|
| ProGen2 | 1440.4 |
| ProteinMPNN | 9.6 |
| ESM-IF | 541.1 |
| Rosetta | 6602.7 |
| DCA | 180.2 |
| LG (Pyvolve) | 5.9 |
| AF-back | 3012.5 |

**Supplementary Table 1**: Runtime using the domain 3CCD:A (50 trajectories, each with 500 accumulated substitutions except for AF-back, where 500 gradient-descent steps were conducted) as an example. The simulation test was performed on the same system equipped with an AMD EPYC 7742 64-core CPU and an NVIDIA A100 (40GB) GPU.

### Sequences of metamorphic pairs

```
>2JP1_A (100% sequence identity)
VGSEVSDKRTCVSLTTQRLPVSRIKTYTITEGSLRAVIFITKRGLKVCADPQATWVRDVV
>1J8I_A
VGSEVSDKRTCVSLTTQRLPVSRIKTYTITEGSLRAVIFITKRGLKVCADPQATWVRDVV

>2QKE_A (95% sequence identity)
MAPLRKTYVLKLYVAGNTPNSVRALKTLNNILEKEFKGVYALKVIDVLKNPQLAEEDKILATPTLAKVLP
PPVRRIIGDLSNREKVLIGLDLLYEEIG
>5JYT_A
MAPLRKTAVLKLYVAGNTPNSVRALKTLANILEKEFKGVYALKVIDVLKNPQLAEEDKILATPTLAKVLP
PPVRRIIGDLSNREKVLIALRLLAEEIG

>2OUG_A (100% sequence identity)
QSWYLLYCKRGQLQRAQEHLERQAVNCLAPMITLEKIVRGKRTAVSEPLFPNYLFVEFDPEVIHTTTINA
TRGVSHFVRFGASPAIVPSAVIHQLSVYKPKDIVDPATPYPGDKVIITEGAFEGFQAIFTEPDGEARSML
LLNLINKEIKHSVKN
>6C6S_D
QSWYLLYCKRGQLQRAQEHLERQAVNCLAPMITLEKIVRGKRTAVSEPLFPNYLFVEFDPEVIHTTTINA
TRGVSHFVRFGASPAIVPSAVIHQLSVYKPKDIVDPATPYPGDKVIITEGAFEGFQAIFTEPDGEARSML
LLNLINKEIKHSVKN

>1UXM_A (99% sequence identity)
```

ATKVVCVLKGDGPVQGIINFEQKESNGPVKVWGSIKGLTEGLHGFHVHEFGDNTAGCTSAGPHFNPLSRK
HGGPKDEERHVGDLGNVTADKDGVADVSIEDSVISLSGDHCIIGRTLVVHEKADDLGKGGNEESTKTGNA
GSRLACGVIGIAQ
>2NAM_A
ATKAVCVLKGDGPVQGIINFEQKESNGPVKVWGSIKGLTEGLHGFHVHEFGDNTAGCTSAGPHFNPLSRK
HGGPKDEERHVGDLGNVTADKDGVADVSIEDSVISLSGDHCIIGRTLVVHEKADDLGKGGNEESTKTGNA
GSRLACGVIGIAQ
